# Model-free Prediction Test with Application to Genomics Data

**DOI:** 10.1101/2022.03.28.486116

**Authors:** Zhanrui Cai, Jing Lei, Kathryn Roeder

## Abstract

Testing the significance of prediction in a regression model is one of the most important topics in statistics. This problem is especially difficult without any parametric assumptions on the data. This paper aims to test the null hypothesis that, given confounding variables *Z*, *X* does not significantly contribute to the prediction of *Y* under the model-free setting, where *X* and *Z* are possibly high dimensional. We propose a general framework that first fits nonparametric regression models on the *Y*|*X* and *Y*|(*X, Z*), then compares the prediction power of the two models. The proposed method allows us to leverage the strength of the most powerful regression algorithms developed from the modern machine learning community. The *p*-value for the test can be easily obtained by permutation. In simulations, we find that the proposed method is more powerful compared to existing methods. The proposed method allows us to draw biologically meaningful conclusions from two gene expression data analyses without strong distributional assumptions: (a) testing prediction power of sequencing RNA for the proteins in CITE-seq data, and (b) identification of spatially variable genes in spatially resolved transcriptomics data.

## 1 Introduction

With the advancement of technology, scientists can collect massive datasets that contain co-variates of interests *X*, confounding variables *Z*, and response *Y* . *X* and *Z* are often high dimensional. A central theme of statistics is to provide modeling and testing tools for the relationship between *X* and *Y* . Traditional statistical theories usually consider parametric models on the data, for example, assuming *Y*|(*X, Z*) follows a normal distribution, or 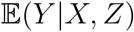 follows a linear model. The modern machine learning community has developed powerful predictive models without parametric assumptions. However, a critical gap remains in the literature: in the model-free setting, how to test whether a set of features have significant predictive power on a response variable. Specifically, we are interested in testing

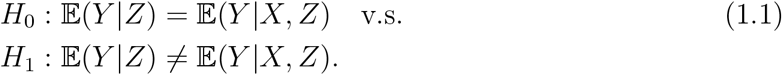

And when only *X* is included in the model, the problem of interest becomes

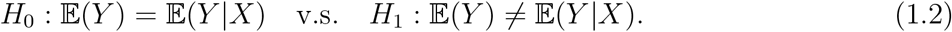

Under the linear model or the single (multiple) index models, the testing problems [1.1] and [1.2] are equivalent to testing whether the coefficient of *X* is equal to zero. From the view of variable selection, [1.1] and [1.2] aim at testing whether *X* is relevant in the prediction of *Y* . Even though the past decades have witnessed many contributions to the statistics literature on variable selection (Fan et al., 2020), it is still extremely challenging to test hypotheses [1.1] and [1.2] without parametric or structural assumptions on *X* and *Y* .

The key idea of our method is to compare whether a powerful machine learning algorithm, fitted with *X* included as part of the input, performs significantly better than without *X*. The method begins by splitting the data into two subsets: *D*_1_ and *D*_2_. We first fit two regression models on *D*_1_: one with *X* and the other without *X*. Then, we compare the performance of the two fitted models on *D*_2_ by calculating the difference between the two means of residual squares. The goal is to detect *any* potential incremental predictive power for *Y* provided by *X* by differentiating the performance of the two models. Under *H*_0_, the two models perform similarly to each other and the residuals should also have similar values. Under *H*_1_, the fitted machine learning algorithm should produce a smaller residual compared to the null model. The test statistic has a limiting normal distribution and the *p*-value can be computed efficiently.

The first application of the proposed method is in CITE-seq data, where surface protein and sequencing RNA are measured simultaneously at the single-cell level. CITE-Seq data is a type of single-cell multimodal omics data, a research area labeled Method of the Year 2019 by Nature Methods. While gene expression data has been extensively studied in the single-cell literature, surface proteins have been less extensively studied because they are relatively difficult to obtain. And yet these proteins are of great interest because they are functionally involved in cell signaling and cell-cell interactions (Davis, 2007). Due to the importance of CITE-seq data in scientific discoveries of human biology, scientists are interested in studying the relationship between proteins and RNA gene expression (Hao et al., 2021; Gayoso et al., 2021). For example, Stuart et al. Stuart et al. (2019) and Hao et al. Hao et al. (2021) used k-nearest neighbors to predict protein levels. Zhou et al. Zhou et al. (2020) implemented deep neural networks to impute surface proteins based on gene expression. Our method can be used to verify whether such models have non-trivial predictive accuracy via a statistically principled test.

The other application arises in spatial transcriptomics data. Scientists have collected high-throughput transcriptome profiling that contains the spatial location of genetic measurements and aim to find the genes that are variable across the tissue. This topic has inspired great interest and Nature Methods recently selected spatially resolved transcriptomics as ‘Method of the Year 2020’ (Marx, 2021). We denote the gene expression as *Y* and the histology information as *X*. Existing literature on spatially variable genes (SVG) detection can be roughly classified into two categories. The first category assumes a parametric model for the distribution of *Y X*. For example, the gene expression profiles *Y* were assumed to follow a normal distribution in (Svensson et al., 2018), given *X* and the spatial structures. The spatial correlation in (Bernstein et al., 2022) is also derived under the normal distribution theory. Because gene expression is usually count data, the other way is to assume *Y* follows a Poisson distribution with rate parameter depending on the spatial correlations among spots (Sun et al., 2020).

The second approach does not assume a parametric model. Some methods utilize certain metrics that measure the spatial distribution within a local radius constraint (Edsgärd et al., 2018; Hu et al., 2020). These method tends to be sensitive to the choice of the local regions in the tissue. Another approach is to test the independence of gene expression and spatial location (Zhu et al., 2021). In comparison, our test specifically target on the expectation of gene expression, and provide great flexibility and interpretative results for the data. We illustrate the details in the numerical analysis.

## 2 Methods

### 2.1 Sample Splitting and Regression

Suppose we observe data (*X*_1_, *Z*_1_, *Y*_1_), . . ., (*X_n_, Z_n_, Y_n_*) independently from the joint distribution of (*X, Z, Y*). We begin by splitting the index 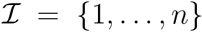 into two subsets 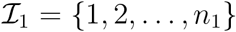 and 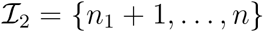. Denote *n*_2_ := *n* − *n*_1_. Let 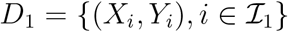 and 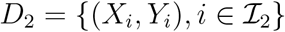 be the two subsets of the data.

The key feature of our method is that it does not rely on a parametric model of 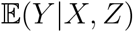, and can easily adapt to different data types, or even high dimensional data. Specifically, we assume the most general form of regression model

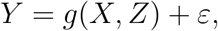

where *ε* satisfies 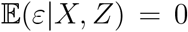. Our method begins with fitting a flexible machine learning algorithm for 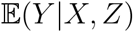 by optimizing

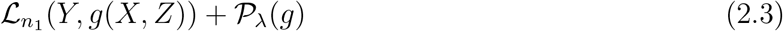

over a function class 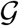 and denote the estimated regression function as 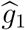. Here ℒ_*n*_1__ (·, ·) is an empirical loss function. When *Y* is continuous, we can choose the least square loss function. We may use the Huber loss when *Y* has heavy tail distribution. When *Y* is discrete, we can use the hinge loss or cross-entropy loss functions. The regularization term *P*_*λ*_ controls the complexity of the estimated model. It is especially useful to let *P*_*λ*_ be the *L*_1_ regularization when the covariates *X* is of high dimension. We also train the model without *X* by optimizing

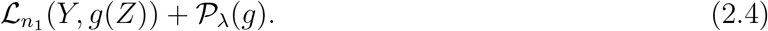

and denote the estimated regression function as 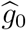.

To ensure the validity of the test, we train 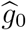 and 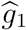 based on the first subset of the data *D*_1_. 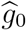 and 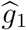 can be fitted using any algorithms, including neural networks, SVM, or random forest. Under *H*_0_, 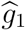 should not perform better than 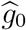 since *X* does not contribute to the prediction of *Y*, while under *H*_1_, a good regression model 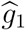 should pick up the information from *X* and result in smaller residuals compared to the null model. This intuition motivates us to implement a two sample test to compare the squared residuals between 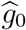 and 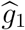 when performing prediction based on *D*_2_.

### 2.2 Two-Sample Comparison

To evaluate the performance of 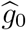 and 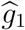, we perform two-sample comparisons on the fitted residuals of the two models based on the data in *D*_2_. The most natural approach is the two sample t-test (TS). Define

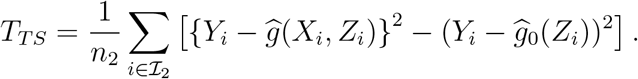

The test will reject *H*_0_ if *T_TS_* takes a large negative value. To give some intuition about the validity of such a test, consider the simpler case where only *X* is included in the model, and the squared loss function is used. In the case 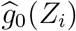 is equivalent to the sample mean of *Y* in *D*_1_, which we denote as 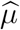.

Under *H*_0_, *X* does not contain predictive information about *Y* so that 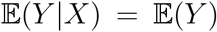 and 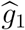 performs no better than 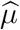, and *T* tends to be non-negative, regardless of the choice of 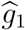. Under *H*_1_, 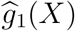 aims to approximate 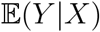, and the positive component in *T_TS_* has a large sample limit of 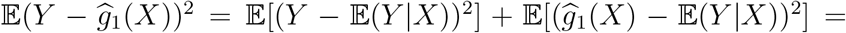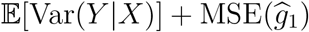. On the other hand, the large sample limit of the negative component in *T_TS_* is 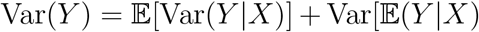. Therefore, when 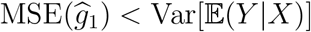, the test statistic *T_TS_* will be negative and the test will have non-trivial power.

To summarize, the test that rejects large negative values of *T_TS_* satisfies the following properties.

1. Under *H*_0_, the false positive is always controlled regardless of the choice of 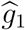.
2. Under *H*_1_, the test has good power as long as 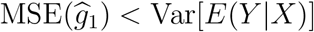.

Our split-fit-test framework allows us to use other forms of two-sample comparisons. For example, in certain scenarios we may want to use the rank-sum test (RS):

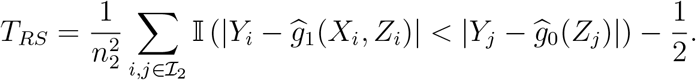

Due to the use of the indicator function, the rank-sum test performs well for data with heavy-tailed distributions or outliers. The intuition behind this test is that when *X* is informative about *Y*, the fitted residuals 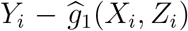 will likely be smaller than those from the null model.

We briefly discuss the pros and cons of the two tests. The rank-sum test, as later shown in the numerical studies, is very robust when the response variable *Y* has a heavy-tailed distribution, but may have inflated type I error if the noise is highly skewed as the quantiles are no longer aligned with expectation. Therefore, we recommend using the rank-sum comparison only if the exploratory analysis does not suggest a highly skewed noise distribution.

In this paper, we obtain the *p*-value using permutation. The algorithm is summarized as follows.

1. For 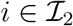, calculate 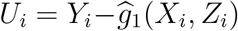 and 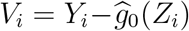. Let 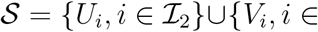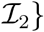.
2. Calculate the test statistic *T* based on *U_i_* and *V_i_*, 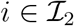.
3. For *b* = 1, *. . ., B*,

a. Obtain sample 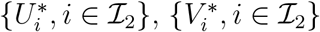 by randomly partitioning *S* into two equalsized subsets.
b. Calculate 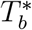 using 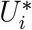 and 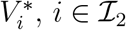.
4. Calculate *p*-value: 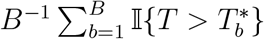.

Under suitable conditions one can show that the test statistic used in the above two-sample residual comparison has a Gaussian asymptotic null distribution. For example, the asymptotic Gaussianity of the *t*-statistic *T_TS_* has been established in Lei (2020). The asymptotic theory for the rank-sum test can be derived using a similar strategy as Cai et al. (2021). As a result, the last step of the *p*-value calculation can be modified by first estimating the standard deviation of 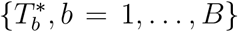, denoted as 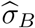, and then calculating the *p*-value as 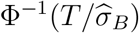, where Φ^−1^ is the cumulative distribution function of standard normal distribution. This will provide a *p*-value with relatively high resolution.

### 2.3 Combine Multiple Splits

The method described so far is based on a single random split of the data. In practice, multiple splits could be used to mitigate the additional randomness introduced by the sample split. To combine the dependent *p*-values obtained from multiple splits, we use the Cauchy combination test proposed by Liu and Xie (2020). The idea is to first transform the *p*-value of each test into a standard Cauchy distribution, then compute the average of the transformed values and compare it with a standard Cauchy’s tail behavior. Specifically, assume we perform *B* splits and obtain *p*-values *p*_1_, . . ., *p_B_*. Let *T*_0_ be defined by

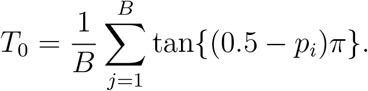

The *p*-value of the combined test can be approximated by

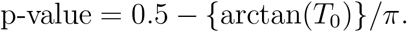

In our numerical studies, we combine the results of 5 – 15 random splits depending on the varying computation cost in different datasets. It turns out that the type-I error of the combined test can be controlled very well and its power exceeds those of single splits.

For notation simplicity, we name the single split regression-based rank-sum test as RS-S, and its multiple version as RS-M. Correspondingly, we name the single split regression-based t-test as TT-S, and its multiple version as TT-M.

## 3 Test Predictability of Proteins in CITE-seq Data

### Background

Cellular Indexing of Transcriptomes and Epitopes by Sequencing (CITE-seq) is a recent multi-modal single cell phenotyping technology (Stuart et al., 2019). The dataset contains measurements of single cell gene expression and surface proteins. Researchers are familiar with gene expression data, which is high-dimensional, noisy, and sparse. By contrast, surface protein data are low-dimensional, highly informative, but more expensive to measure. Thus, it is of great interest to predict protein measurements based on gene expression Stuart et al. (2019); Zhou et al. (2020); Gayoso et al. (2021); Hao et al. (2021). These prediction models provide a better understanding of the translation from RNA-seq to proteins and also enable researchers to predict the proteins when only the RNA sequence is measured at the single cell level.

We will use the human PBMC CITE-seq data, which has been analyzed in Hao et al. (2021), as our primary example. While different types of cells usually contain different patterns of gene expression and proteins, it is unclear how the predictability of proteins varies across cell types. In this section, we will investigate the predictability of protein expression in different types of human blood immune cells.

### 3.1 Simulations

To verify the performance of the proposed method, we perform simulations for the predictive tests based on rank-sum test and two sample t-test. We consider both single split and multiple split of the data, and set the split times in multiple split to be 10. XGBoost tree (Chen and Guestrin, 2016) is implemented as the regression algorithm due to its fast computational speed and good flexibility to capture nonlinear relationships. We compare our method with the martingale difference correlation (MDC), which has the similar goal of testing mean independence and was proposed by Shao and Zhang in (Shao and Zhang, 2014).

To demonstrate the performance for high-dimensional, sparse signal and heavy-tail noise, we generate the response *Y_i_* as the function of the first element of the covariates *X*_*i*,1_ with a heavy-tail distributed error term. Specifically, let

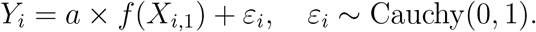

*a* is used to control the signal level. *a* = 0 implies that *H*_0_ is true and *a* > 0 represent *H*_1_ holds. The details of *f*(·) in each model is given in the Supplementary Information.

We consider two simulation scenarios: 1) the signal level *a* increase from 0 to 1 when sample size and dimension are fixed to be 200 and 1000. 2) the dimension of *X* increase from 100 to 1000 when keeping the sample size and *a* fixed. Each simulation is repeated 1000 times and we report the average power in Fig.1 and 2, where the type-I error is controlled at *α* = 0.05. As we can see, all methods have a valid type-I error rate. The multiple split rank-sum test performs the best in terms of power, followed by the single split rank-sum test. The two sample t-test seems to be unsuitable for the heavy-tailed data. As for MDC, we see that it not only is unsuitable for heavy-tailed data but also suffers from the curse of dimensionality when the covariates is high-dimensional. This demonstrates the advantage of the proposed methods.

**Figure 1:**
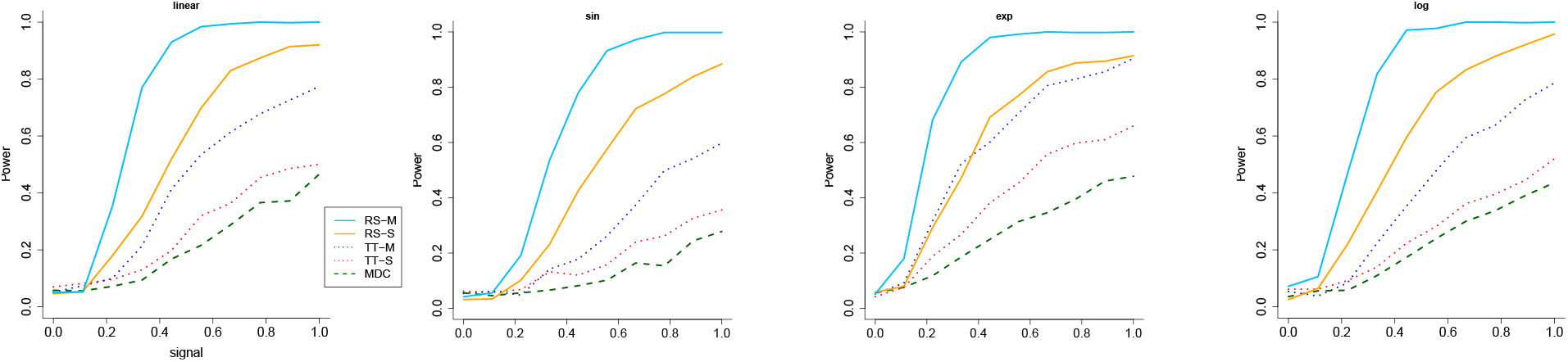
The power (vertical) versus signal (horizontal) for the all the methods when the response has heavy-tailed distribution and *α* = 0.05. The RS stands for rank-sum test and TT stands for two sample t-test. M means multiple split and S represents a single split.

**Figure 2:**
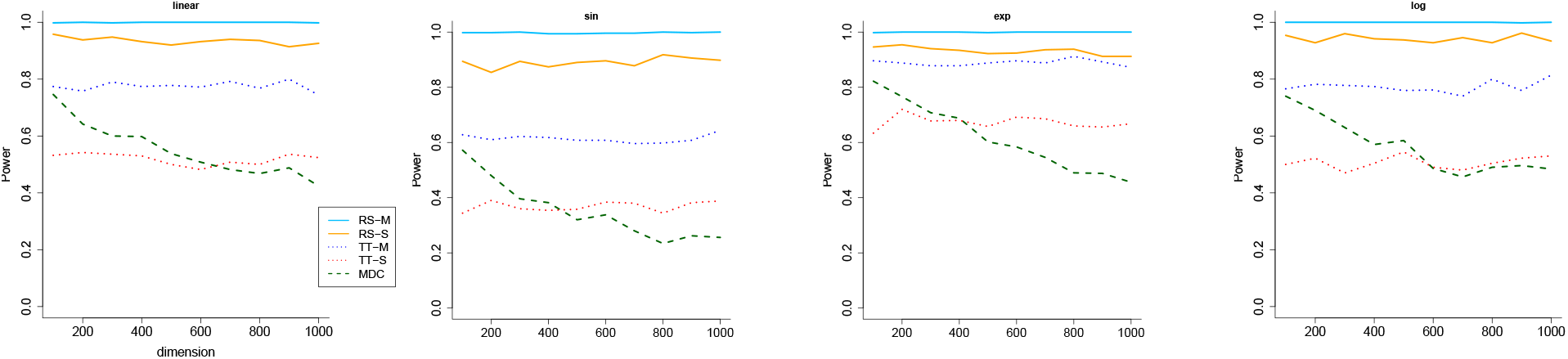
The power (vertical) versus dimension (horizontal) for the all the methods when the response has heavy-tailed distribution and the dimension of the covariates increases. *α* = 0.05. The RS stands for rank-sum test and TT stands for two sample t-test. M means multiple split and S represents single split.

### 3.2 Human PBMC Data

After applying standard quality control procedures Hao et al. (2021), the human PBMC data contains 20,729 gene expressions and 228 proteins measured on 161,764 cells. After removing the 2 proteins (CD26-1 and TSLPR) that contain mostly zeros, we obtain 226 proteins in total. According to the cell annotations, we can classify all the cells into 8 different types. Following the standard convention in single cell analysis, we restrict our analysis to the top 5,000 highly variable gene set. The marker genes for each cell type are obtained by implementing the *FindMarkers* function from Seurat (Hao et al., 2021). The number of cells and marker genes of each type are summarized in Table 1. The total number of marker genes is less than the sum of individual cell types, because different cell types may share the same marker genes. Under the testing framework of [1.1] and [1.2] We are mainly interested in two questions:

1. Do marker genes alone provide prediction power on proteins?
2. Besides marker genes, do other gene clusters provide additional prediction power for proteins?

**Table 1:** Number of cells and marker genes in each cell type in the human PBMC data.

| Cell type | Cells | Marker genes |
| --- | --- | --- |
| Mono | 49,010 | 1,007 |
| CD4 T | 41,001 | 576 |
| CD8 T | 25,469 | 313 |
| NK | 18,664 | 519 |
| B | 13,800 | 598 |
| other T | 6,789 | 122 |
| DC | 3,589 | 372 |
| other | 3,442 | 273 |
| Total | 161,764 | 1,692 |

**Table 2:** Cell type and protein names where the top 5,000 genes reject *H*_0_ but the marker genes fail to reject

| cell type | protein |
| --- | --- |
| CD8 T | TCR-2 |
| other | Siglec-8 |
| Mono | CD284 |
| CD8 T | CD49a |
| CD4 T | TCR-V-7.2 |
| CD4 T | CD177 |
| other | CD8a |
| other | CD324 |
| NK | CLEC2 |
| other | CD3-2 |

To answer these two questions, we implement our method by using XGBoost tree and the rank-sum test with 5 splits. In each cell type, we test the predictability of both the top 5000 highly variable genes and the marker genes. For each protein, we treat the gene transcription as *X* and protein as *Y* in testing [1.2]. Because 226 tests are conducted for each cell type, we adjust all the p-values by applying the Bonferroni correction method to control the familywise error rate at 5%.

To better illustrate the similarity of the testing results for the top 5,000 genes versus the marker genes, we display the testing results row by row in Fig.3: specifically, the odd and even rows correspond to the predictability of the top 5,000 genes and the marker genes, respectively. The columns represent the 226 proteins. Blue and orange bars highlight the proteins/cell type combinations for which we reject *H*_0_. Notably the testing results for the two batches of genes are quite similar to each other: more than 97% of the tests reach the same conclusion. For the tests that disagree, 0.6% of the tests are the cases where the top 5,000 genes reject *H*_0_ but the marker genes fail to reject. Those cases cover 10 proteins with its cell type given in SI Table.2. For those cases, the interaction effects between the marker genes and other genes might provide additional prediction power. Besides, 2.3% of the tests are the cases where the marker genes reject *H*_0_ but top 5,000 genes fail to reject. We believe this is because increased noise in the data lead to inferior performance of the regression algorithm. It is also interesting to observe that among the 226 proteins, 17 proteins can be predicted in all cell types and 36 proteins fail to be predicted in all cell types, based on the inference results using the top 5,000 genes. The marker genes tell us the same story. It is interesting to find that the mean value for the 36 proteins is 1.38 while it is 12.63 for the 17 proteins. In general, proteins with rich measurements can be predicted well.

**Figure 3:**
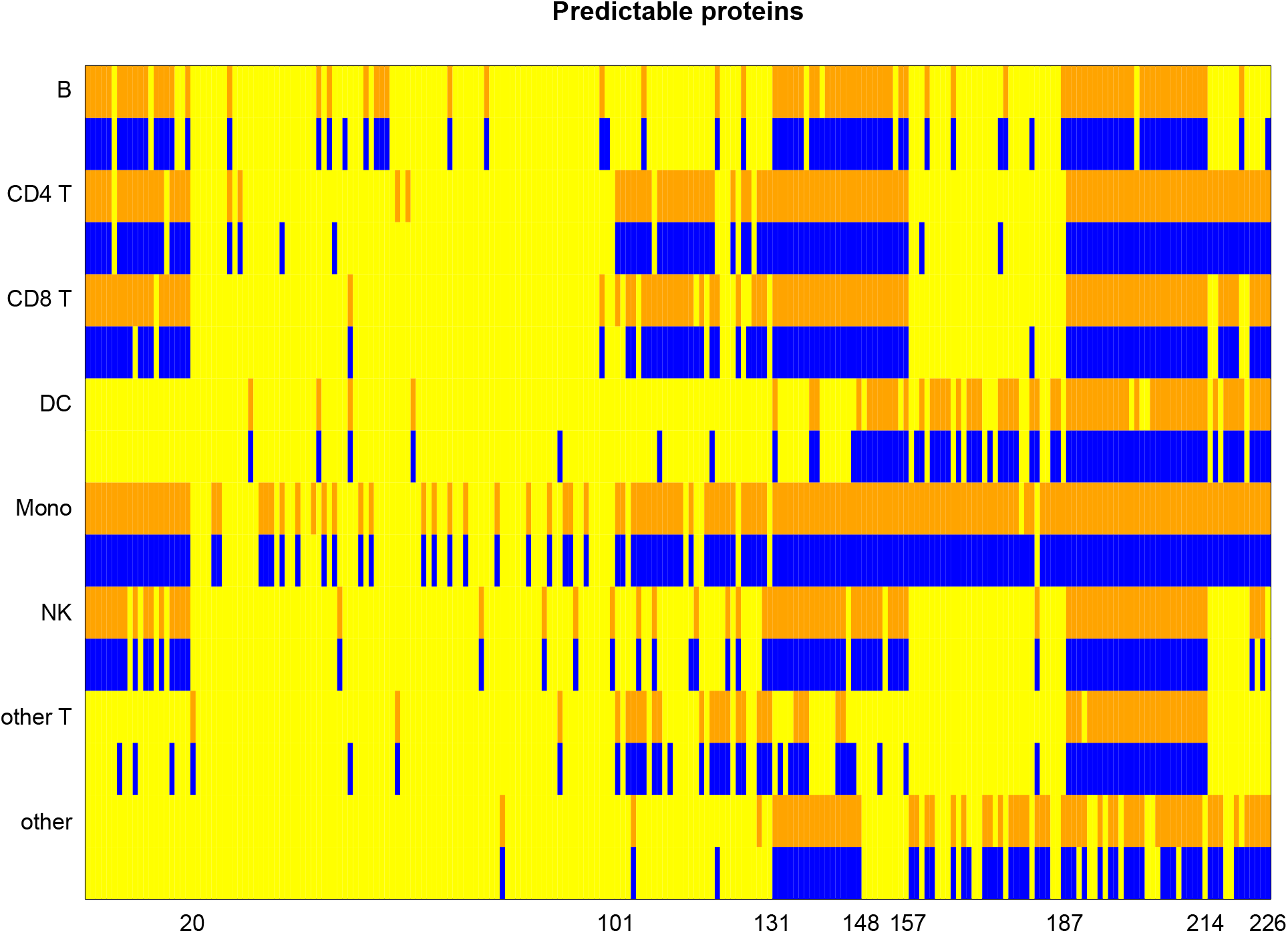
The predictability test of every protein in all cell types. The columns represent the 226 proteins, and the rows represent the 8 different cell types. Under the testing problem [1.2], the orange rows represent *X* being all 5000 genes, while the blue rows represent *X* being the marker genes.

To answer the second question, we let 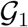 denote the marker genes for each cell type. Then we remove the 1,692 marker genes from the set of top 5000 genes and perform clustering analysis (Morabito et al., 2021) for the remaining genes. We obtain 13 clusters, denoted as 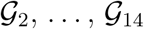, the size of which decrease from hundreds to dozens. The goal is to test the prediction power of 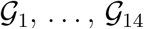 on the proteins in each cell type.

We begin by testing the prediction power of the marker genes 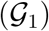 in each cell type. Specifically, each protein is treated as *Y* and 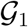 is treated as *Z* in the testing problem [1.1]. Then, we test whether adding 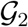, . . ., or 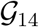 to 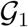 improves the predictability of each protein. This is achieved by treating each of 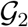, . . ., or 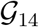 as *X* in testing problem [1.1]. We present the inference results for NK cells in Fig.4, and relegate the other cell types to Figure29-35 of the Supplementary Information, where the p-values are also adjusted by Bonferroni correction. Similarly, the blue bars represent the rejection of *H*_0_ while the yellow bars represent the failure of rejection. The rows correspond to 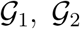, . . ., etc, and the columns represent the proteins. As we expect, the marker genes are extremely useful in predicting the proteins. Adding extra gene clusters occasionally provides extra prediction power, and one protein (CD177) that is not predictable by marker genes is predictable with another gene cluser.

**Figure 4:**
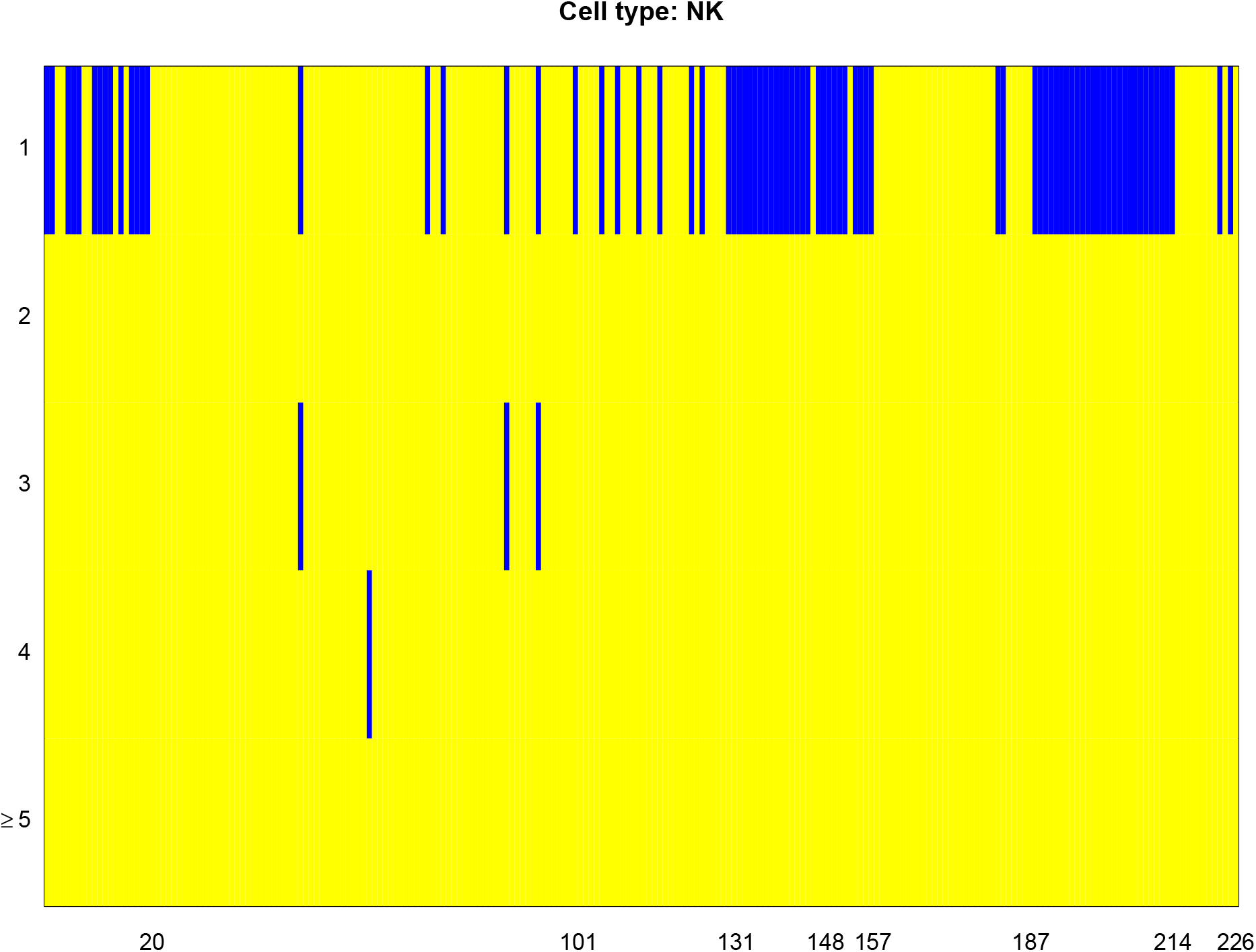
The predictability test of every proteins for NK cells. The first row represents the group marker genes 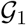, and the second to the last rows represent the other clusters of genes 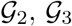,.... We combine several clusters in the last row since the rejection of *H*_0_ becomes rare.

## 4 Spatially Variable Genes Detection

### 4.1 Background

Recent technological advances in spatially resolved transcriptomics have enabled gene expression profiling with spatial information on tissues. The spatial transcriptomics sequencing technique measures the expression level for thousands of genes in different spots, which may contain multiple cells. The smFISH technique detects several mRNA transcripts simultaneously at the sub-cellular resolution but usually has relatively low expression levels compared to spatial transcriptomics. More recent technologies such as MERFISH (Moffitt et al., 2018) and seqFISH (Shah et al., 2016) can substantially increase the number of detectable mRNAs from hundreds to thousands.

The gene expression data is represented in an *n* × *p* matrix, where each column denotes a specific gene, and each row denotes an observed sample spot in the tissue. Each spot may contain one or multiple cells depending on the experimental method. The spot is also associated with a 2-dimensional spatial location in the sample. It is of interest to identify the genes that display spatially distinct expression patterns. Similar to existing approaches (Sun et al., 2020; Svensson et al., 2018), we first test each gene separately, then control the false discovery rate based on the *p*-values for all genes. For each gene, we denote its expression level as *Y* = (*Y*_1_, *, Y_n_*), and the corresponding spatial location at the *n* spots as *X* = (*X*_1_, *. . ., X_n_*). In our numerical analysis, we let *X_i_* be a four-dimensional covariate: intercept, horizontal coordinate, vertical coordinate, and the interaction effect of horizontal and vertical coordinate. Under the testing framework [1.2], we are interested in finding the genes that satisfy 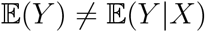.

### 4.2 Simulations

We aim to determine via simulations if two versions of the proposed method (TT-S and TT-M) work well when compared with popular SVG methods, SPARK (Sun et al., 2020) and SpaDE (Svensson et al., 2018). Both methods are parametric models: SPARK assumes the gene expression follows Poisson distribution and SpaDE assumes the data follows Gaussian distribution.

To generate synthetic data, we use the spatial location of the upcoming mouse olfactory bulb data and generate our spatial signals. Following Sun et al. (2020), we consider three spatial patterns: hot spot, gradient, and streak. A generic picture of all the three simulated models is illustrated in Figure 5. The random forest algorithm is implemented in the regression step. We generate the signals according to the three patterns, and add a random noise that follows the uniform distribution on [0, 1] to each spot. Specifically, denote the signal as *f* (*X_i_*), where *X_i_* is the spatial information at location *i*. The gene expression at location *i* is generated by

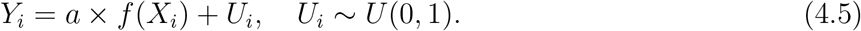

**Figure 5:**
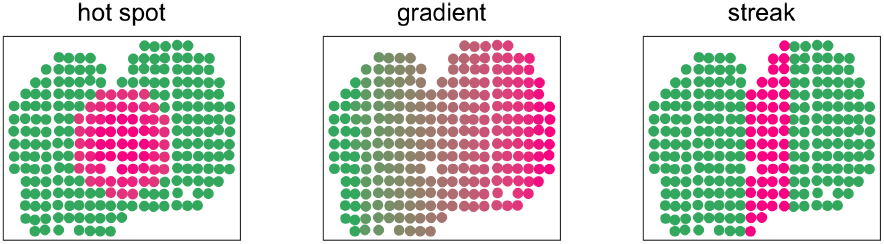
The three spatial patterns for the gene expression in the simulation settings.

The details of *f*(·) for each spatial pattern is given in the Supplementary Information. The signal level is controlled by the constant *a* ∈ [0, 1] in equation (4.5), where *a* = 0 implies the *H*_0_ is true and *a* > 0 indicates that *H*_1_ holds. Because the real data are sparse, we also consider the settings where we set *Y_i_* = 0 if *Y_i_* < median(*Y*). Thus 50% of all the locations are set to zero.

The simulation is repeated 1000 times and we report the average power for all the methods. The random forest algorithm is implemented as the regression tool and we set the number of multiple splits to be 15. The power curves are illustrated in Figure 6 where the type-I error is set at 0.05. Additional simulation results are reported in the Supplementary Information. The *Y*-axis is the power and the *X*-axis is the signal level *a*. When *a* = 0, the null condition holds, and all methods can control the type-I error very well. As *a* increases, the power also increases as expected. The proposed method shows superior performance across all settings: hot spot, gradient, and streak. The power curves also show that multiple splits are indeed more powerful compared to a single split under the alternative hypothesis. As the comparison between the two rows, we also find that all methods tend to perform better when the data are sparse, where half of the low expression levels are set to 0. This might be due to the low noise level under the sparse setting. SPARK tends to perform better than SpaDE under the hot spot and streak settings, and SpaDE outperforms SPARK when the signal is gradient. This finding also echos the simulation results in Sun et al. (2020) where SPARK and SpaDE have similar power only when the signal is gradient.

**Figure 6:**
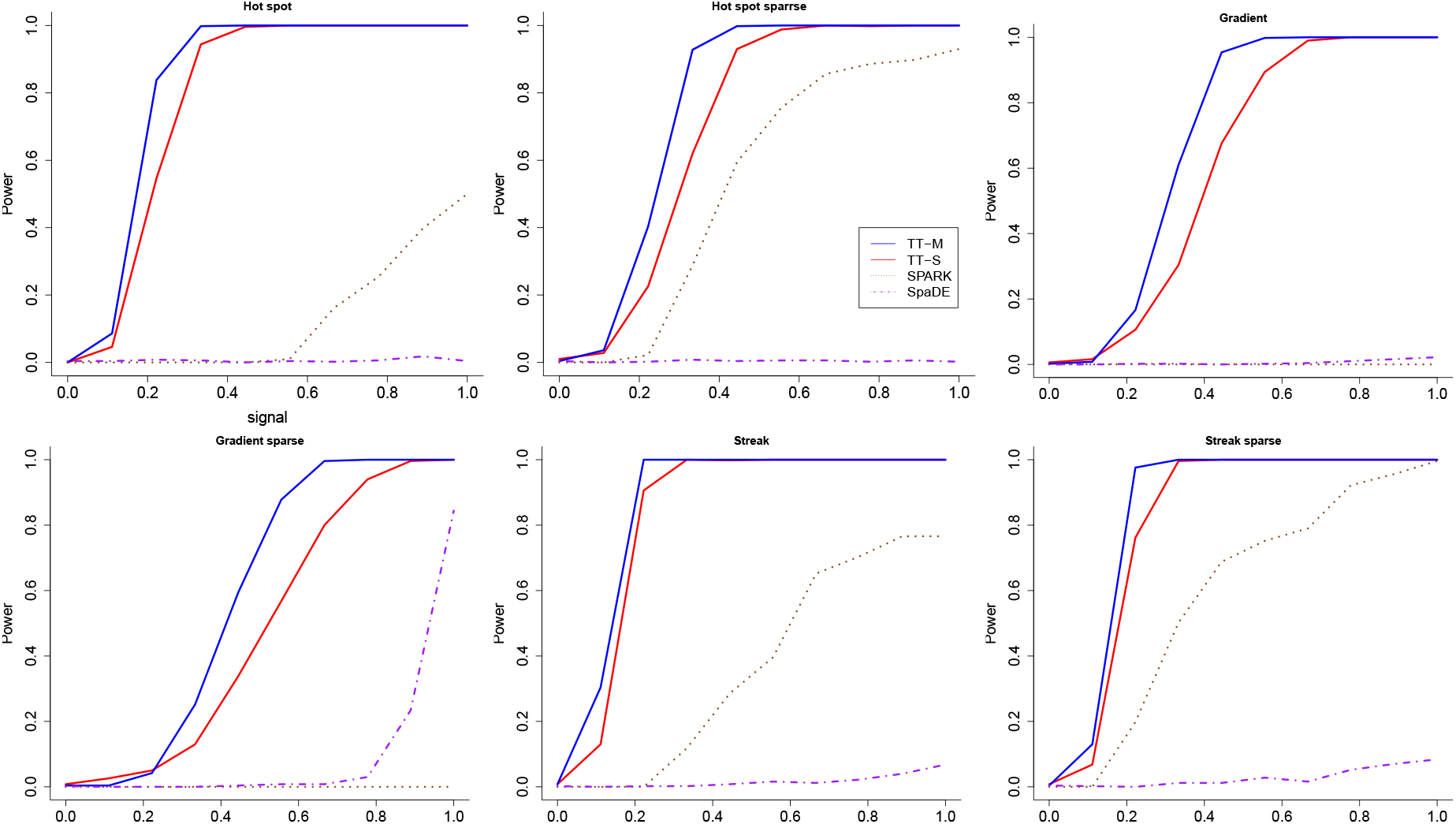
The power (vertical) versus signal (horizontal) for the regression-based two sample t-test, SPARK, and SpaDE when *α* = 0.05. The three rows correspond to the three patterns: hotspot, gradient and streak. The two columns represent non-sparse and sparse settings.

Next, we apply the proposed test to real datasets. The analyses illustrate that our proposed method produces calibrated *p*-values under the null condition in the randomly permuted data, and shows impressive power compared with existing approaches. We advocate the validity of our results since they do not require any distributional assumptions on the gene expression data and have less constraint when applied to real data. Because the real data analysis involves multiple testing, we adjust all the *p*-values by applying the Benjamini-Yekutieli method Benjamini and Yekutieli (2001) to control the false discovery rate at 5%. The choice of regression algorithm and the number of multiple splits are set to be the same as in the simulation.

### 4.3 Mouse Olfactory Bulb Data

Spatial transcriptomics sequencing was used to produce the mouse olfactory bulb data (Ståahl et al., 2016). Following previous analyses using SpaDE (Svensson et al., 2018) and SPARK (Sun et al., 2020), we used the MOB Replicate 11 file, which contains 16,218 genes measured on 262 spots. Similar to Sun et al. (2020), we filter out genes that are expressed in less than 10% of the array spots and select spots with at least ten total read counts. After the filtering, we obtain 11274 genes on 260 spots.

All methods produce valid *p*-values under the null condition where the response is randomly permuted (Fig.7a). With the original data, SPARK, SpaDE, TT-S, and TT-M identified 772, 68, 234, and 731 spatially variable genes, respectively. More than 40% of the genes identified by TT-M overlap with SPARK (Fig.7b). For the Upset plot, the left bar plot represents the total size of each set, and the top bar plot represents the intersection of each methods. Every possible intersection is shown by the bottom plot.

**Figure 7:**
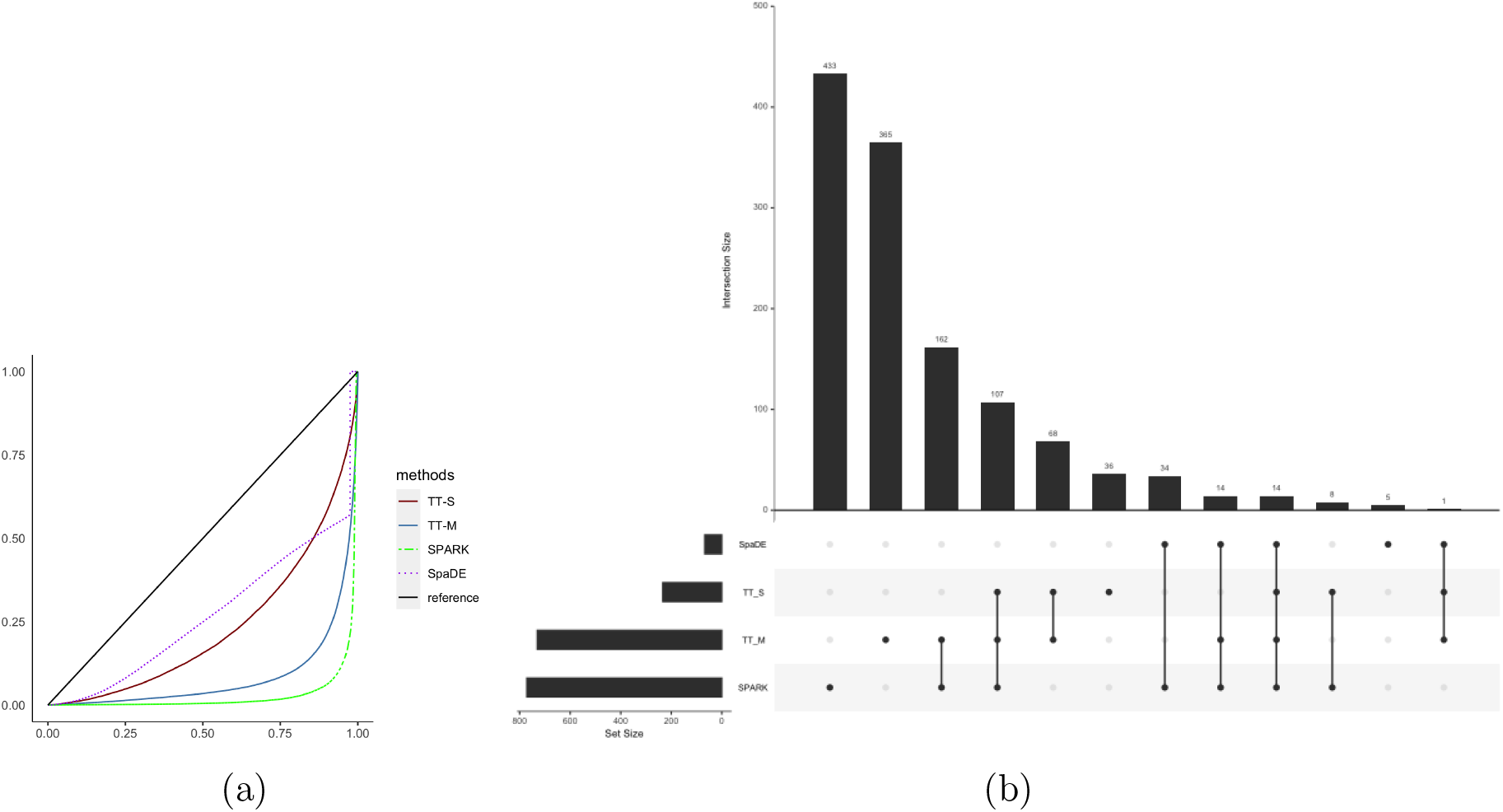
Analysis for the mouse olfactory bulb dataset. (a): The empirical distribution of the *p*-values under the null condition in the permuted data. The blue solid line and red solid line denote the multiple splits (TT-M) and single splits (TT-S). The green dashed line denotes SPARK, and the purple dotted line denotes SpaDE, respectively. (b): The Upset plot shows the overlap of genes for all the four methods. The left bar plot represents the total size of each set, and the top bar plot represents the intersection of each methods. Every possible intersection is shown by the bottom plot.

As an advantage of our method, we can check the variable importance of each feature as part of the random forest algorithm. Specifically %IncMSE gives the increase in mean square prediction error as a result of the target variable being randomly permuted, and a larger value indicates relatively higher importance. In our data analysis, the average %IncMSE is 0.072 for the horizontal axis, 0.044 for the vertical axis, and 0.056 for the interaction effect of the horizontal and vertical axis. This indicates that most spatial variability in expression occurs across the horizontal axis as depicted by the most significant 8 genes (Fig.8(a)). The variation in expression is notable and approximately symmetric in the horizontal axis indicating that the proposed method captures the variability in the spatial distribution accurately.

**Figure 8:**
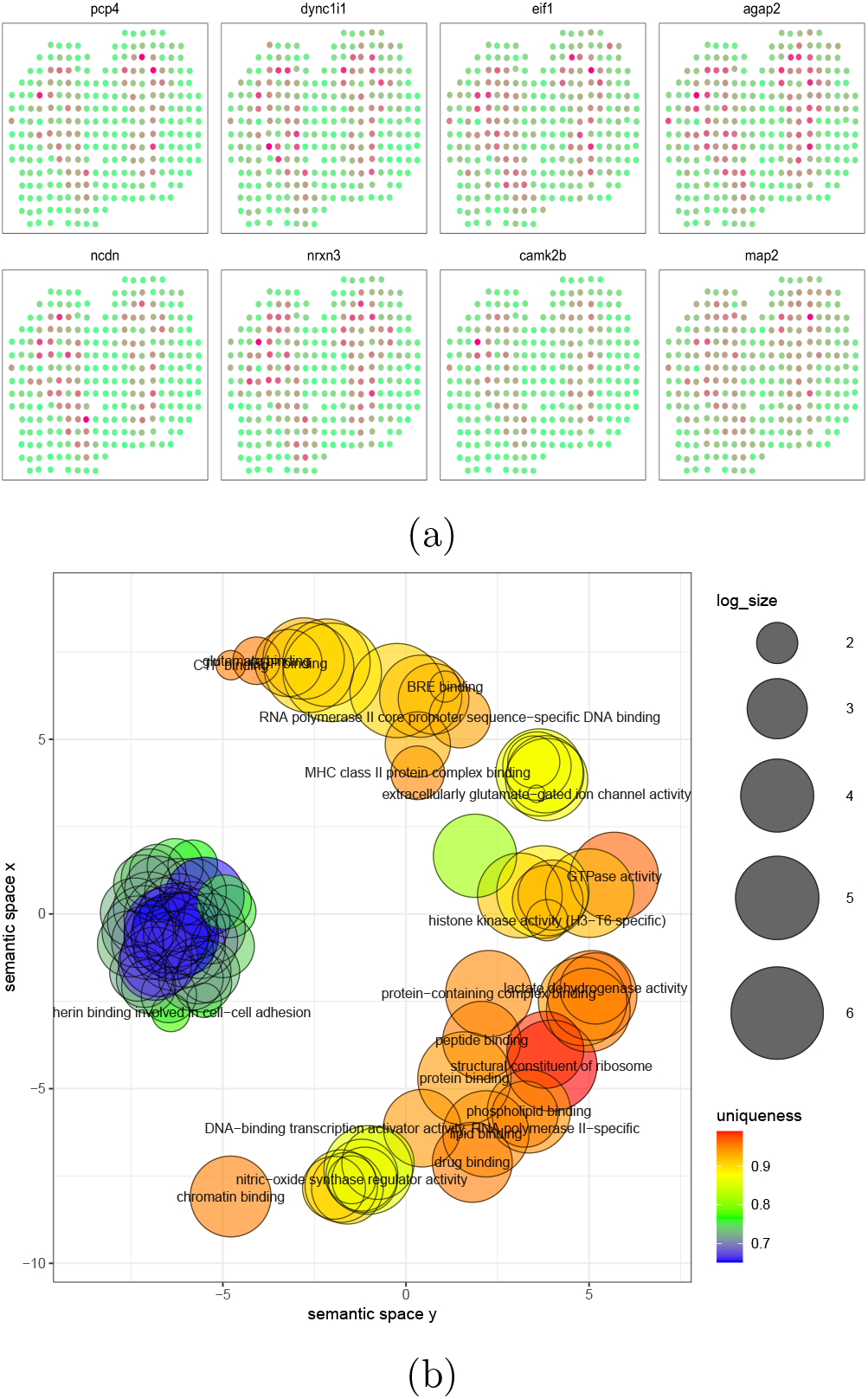
Analysis for the mouse olfactory bulb dataset.(a): The 8 genes that have the smallest *p*-values detected by TT-M. (b): The clustering of GO annotations for the genes detected by TT-M.

Lastly, we perform GO enrichment analysis for molecular function and display the clustered GO annotations by implementing Revigo Supek et al. (2011). The enrichment results offer an understanding of the detected spatially variable genes detected by our method (Fig.8b, SI, Fig.17 and 18). Most of the detected genes are related to the binding of certain proteins or DNA. For example, the cluster in the left colored in blue and green represent the genes that are essential to cadherin binding in cell-cell adhesion. Our results complement the GO terms identified by SPARK which are related to synaptic organization and olfactory bulb development.

### 4.4 Human Breast Cancer Data

The human breast cancer data are also obtained by spatial transcriptomics sequencing (Ståahl et al., 2016). Following previous analyses using SpaDE (Svensson et al., 2018) and SPARK (Sun et al., 2020), we use the Breast Cancer Layer 2 file, which contains 14,789 genes measured on 251 spots. We filter out the genes that are expressed in less than 10% of the array spots and selected spots with at least ten total read counts. After filtering, we obtain 5262 genes measured on 250 spots.

The results are summarized in Fig.9. As expected, all methods produce valid *p*-values under the null condition. SPARK identified 290 and SpaDE identified 115 genes spatially variable genes. By comparison TT-S identified 335 genes with more than 1/3 overlaping with SPARK and TT-M identified 701 genes with around 1/4 overlapping with SPARK. Our proposed methods found considerably more genes compared to existing methods. We found that the breast cancer data has 23% of non-zero elements in the gene expression matrix while the mouse olfactory bulb data has 56% of non-zero values. One possible explanation is that our methods are more powerful in picking up weak signals in sparse gene expression data.

**Figure 9:**
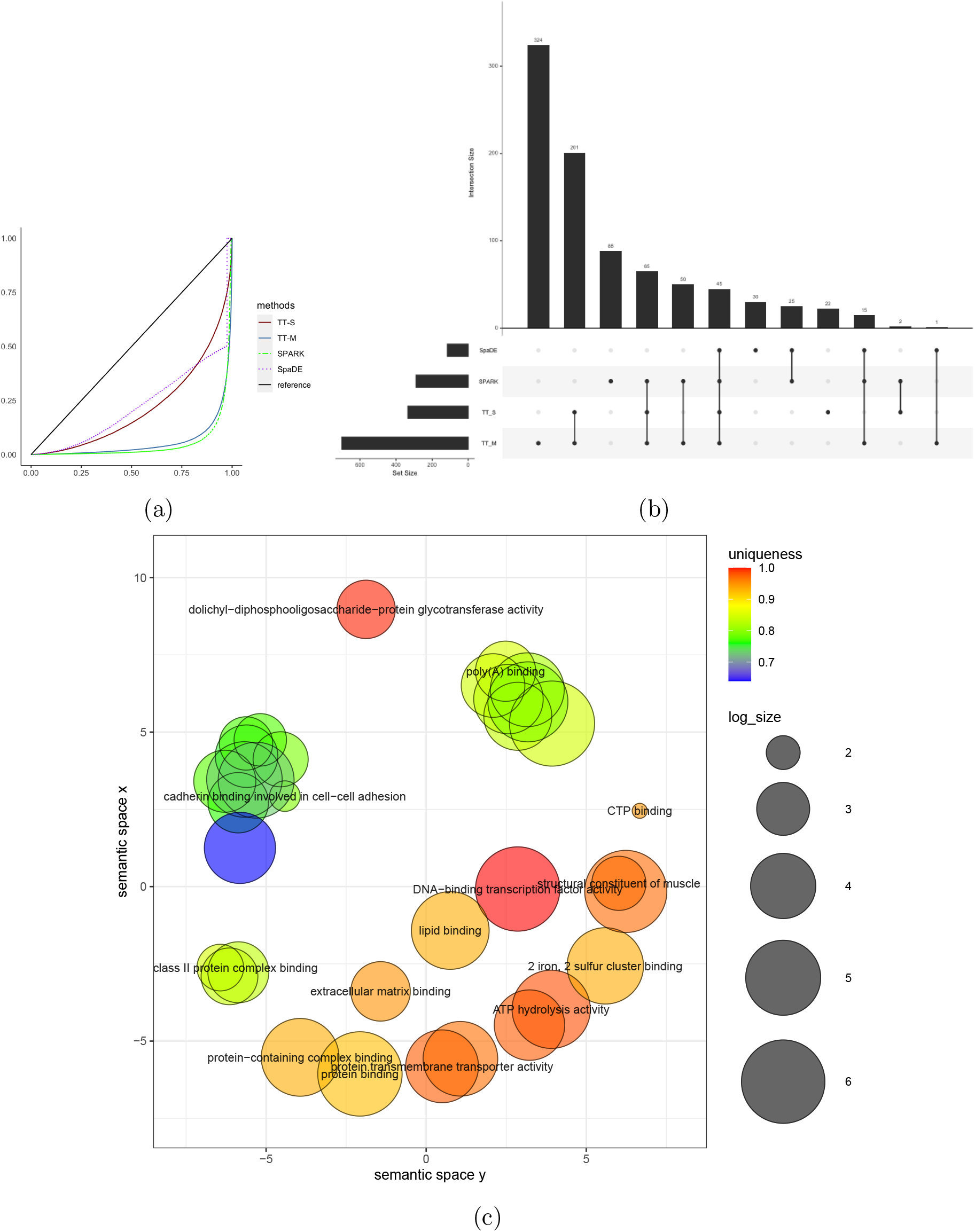
Analysis for the human breast cancer data. (a): The empirical distribution of the *p*-values under the null condition in the permuted data. The blue solid line and red solid line denote the multiple splits (TT-M) and single splits (TT-S). The green dashed line denote SPARK, and the purple dotted line denotes SpaDE, respectively. (b): The Upset plot shows the overlap of genes for all the four methods. (c): The clustering of GO annotations for the genes detected by TT-M.

To provide additional evidence of the findings, we look into the overlaps of the detected genes with background information. We found 8 among the 14 cancer-related genes that are highlighted in the original study (Ståahl et al., 2016). SpaDE detected 7, SPARK detected 9, see Fig.10c for the overlaps of those genes. The gene expression of the 8 detected genes are illustrated in Fig.10a and the 6 missed genes are plotted in Fig.10b. Clearly, the detected genes show strong spatial patterns. We also found 79 genes that are previously known to be related to cancer according to the CancerMine database (Lever et al., 2019). On the other hand, SpaDE detected 11 and SPARK detected 40.

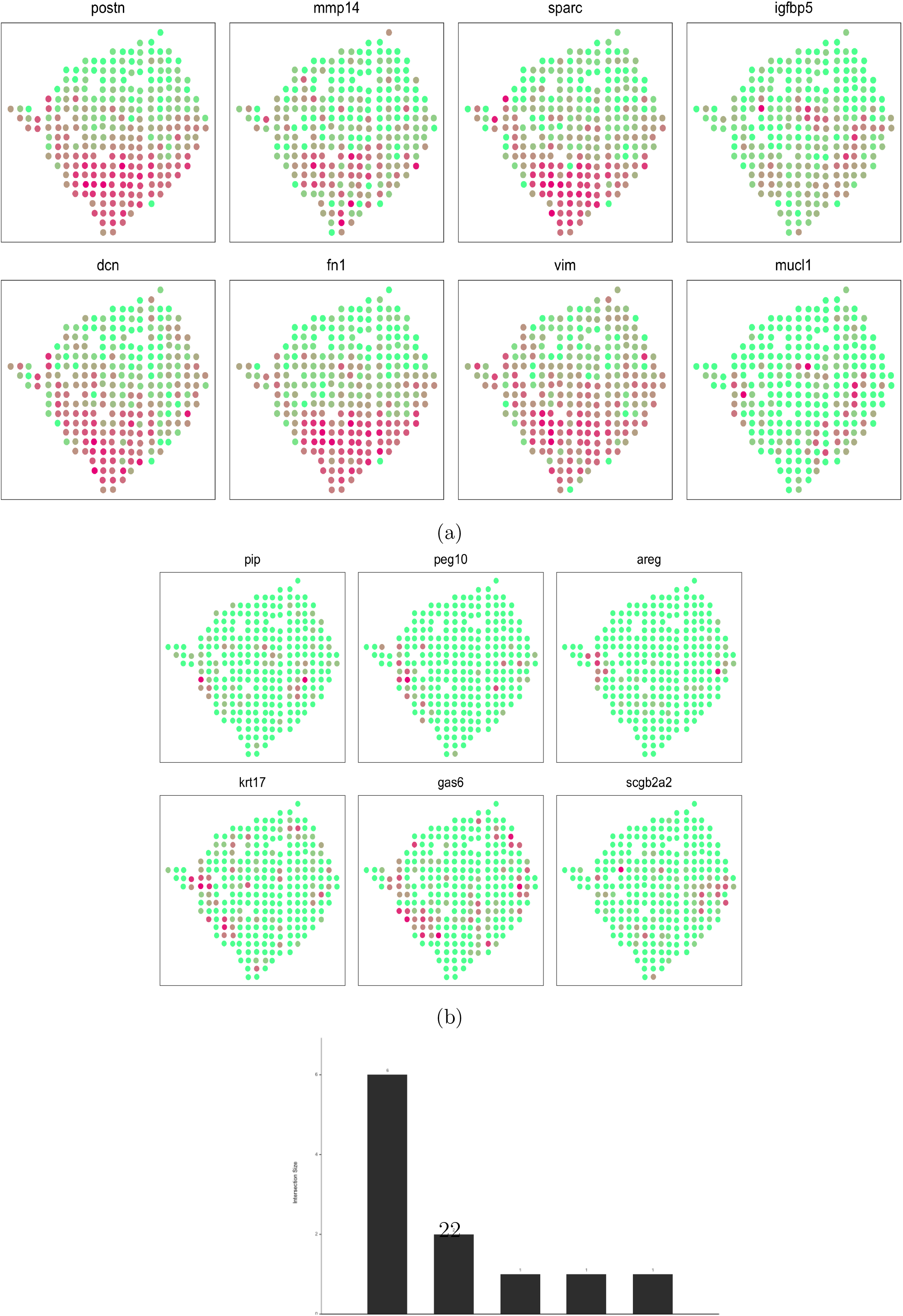

In terms of variable importance, the average %IncMSE equals 0.036 for the horizontal axis, 0.120 for the vertical axis, and is 0.070 for the interaction effect of the horizontal and vertical axis. In contrast to the analysis of the mouse olfactory bulb data (Fig.8a), the vertical axis plays a more important role in the spatial patterns. This phenomenon can be observed in the detected cancer genes shown in Fig10a. The enrichment results provide deep understanding of the detected spatially variable genes (Fig.9c, Supplementary Information Fig.19 and 20). Most of the detected genes are also related to bindings of important cell functions and proteins.

## 5 Discussion

In this paper, we proposed a new approach for the test of covariates and applied the new method to both test of prediction power in CITE-seq data and the identification of spatially variable genes. Distinguished from the previous methods, the proposed method does not assume any parametric distributions on the gene expression data, which provides great flexibility for real data analysis. We are also able to implement a large class of machine learning regression algorithms in the test, such as neural networks, random forest, SVM, etc.

Due to the sample splitting and machine learning algorithms, our test may not perform well when the sample size is too small. In the analysis of a small seqFIISH data shown in the supplementary information, the proposed test found a relatively small number of spatially variable genes. The sample size in this data is only 131. Thus the effective sample size for training the random forest is only 65, which is not enough to obtain a properly trained algorithm.

There are several potential works left for future research. For CITE-seq data, both the dimension and sample size are huge, and the design matrix is extremely sparse. This type of data presents unique challenges and its analysis requires further development of theoretical and computational statistical tools. In spatial transcriptomic studies, the current literature on the test of spatially variable genes is all based on single tests applied on each gene in the domain. The control of false discovery rate is achieved by simply applying either qvalue Storey (2003) or the Benjamini-Yekutieli method Benjamini and Yekutieli (2001). How to incorporate spatial information to achieve better false discovery control performance is a very promising future research topic.

### A Details of Simulations in Predictability Test

We let sample size *n* = 200. *X* ∈ ℝ^1000^ is generated from the standard normal distribution and *ε* ∈ ℝ^1^ is drawn from the standard Cauchy distribution. *Y* is set to be only related to *X*_1_ (the first element in *X*) and *ε*. We consider very simple models, as those are most effective in illustrating the power performance of all methods.

- *Y* = *a* × 10*X*_1_ + *ε*;
- *Y* = *a* × 10 sin(*X*_1_) + *ε*;
- *Y* = *a* × 10 exp(*X*_1_) + *ε*;
- *Y* = *a* × 100 log(*X*_1_ + 10) + *ε*;

The signal level *a* is set to vary from 0 to 1 to fully investigate the size and power of each test. We repeat the simulation 1000 times and report the average results. We report the additional power curves for all the methods in Fig. 11 and 12, where the type-I error rate is set as 0.01. Our methods perform very well under all settings. Moreover, the increasing dimension does not have negative effects on the proposed methods. For MDC, the power decreases dramatically as the dimension of *X* increases from 100 to 1000.

**Figure 11:**
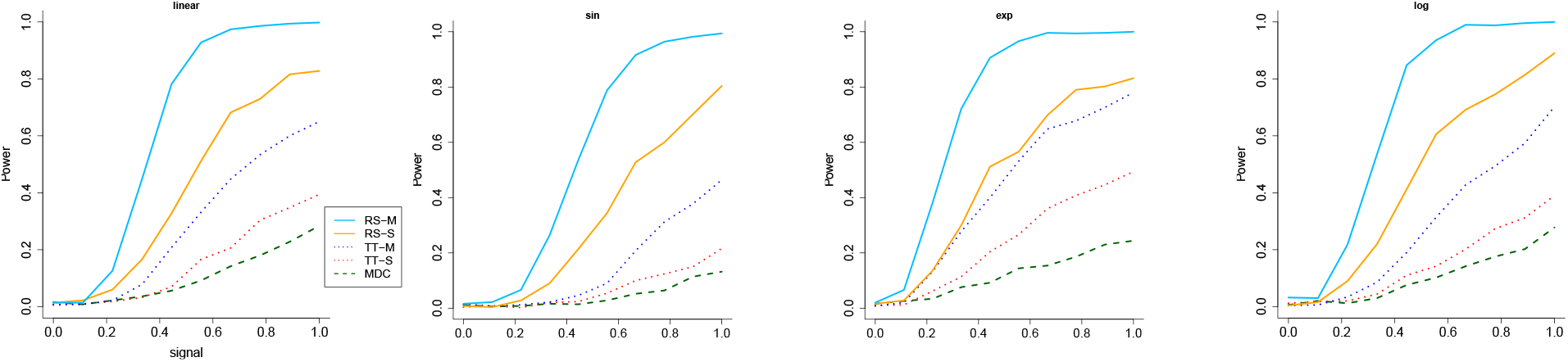
The power versus signal for the all the methods when the response has heavy-tailed distribution and *α* = 0.01. The RS stands for rank-sum test and TS stands for two sample t-test. M means multiple split while S represents single split.

**Figure 12:**
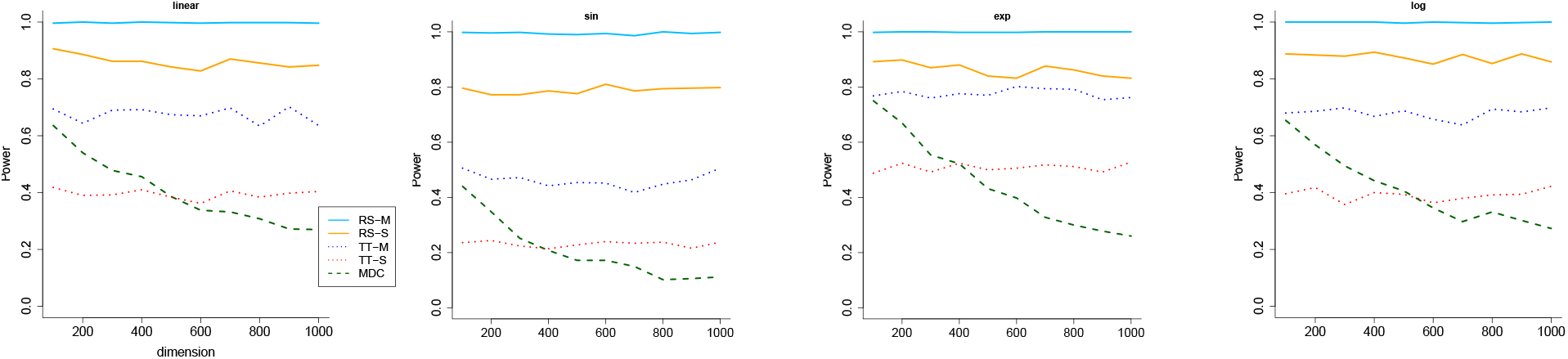
The power versus dimension for the all the methods when the response has heavy-tailed distribution and the dimension of the covariates increases. *α* = 0.01. The RS stands for rank-sum test and TS stands for two sample t-test. M means multiple split while S represents single split.

### B Details of Simulations in Spatially Variable Genes Detection

To compare the power of each test under both null hypothesis and alternative hypothesis, we generate the signals according to patterns, and add a random noise that follows uniform distribution on [0, 1] to each spot. Specifically, denote the signal as *f* (*X_i_*), where *X_i_* is the spatial information at location *i*. The gene expression at location *i* is generated by

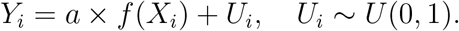

For all three patterns, we describe the model details as follows.

#### Hotspot

We randomly choose a spot 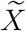 whose horizontal and vertical axis both follow a uniform distribution between the range of {*X*_1_, . . ., *X_n_*}. Let *d_i_* denote the Euclidean distance between *X_i_* and 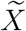, then we set

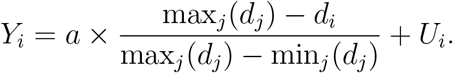

#### Gradient

Let *X*_*i*,1_ denote the horizontal axis of spot *i* and *X_m_* be the smallest horizontal axis, i.e., *X_m_* = min_*j*_(*X*_*j*,1_). Then we set

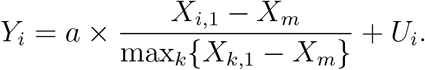

#### Streak

We randomly choose a spot 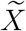 whose horizontal axis is between the 0.4 and 0.6 quantile of the horizontal axis of {*X*_1_, . . .,*X_n_*}. Let *h* be a tuning parameter adjusting the width of the streak. Then

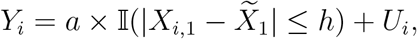

We repeat the simulation 1000 times and report the average results. Besides the figures in the main paper, we also report the power performance of all methods in Figure 13 where the type-I error *α* is set to be 0.01.

**Figure 13:**
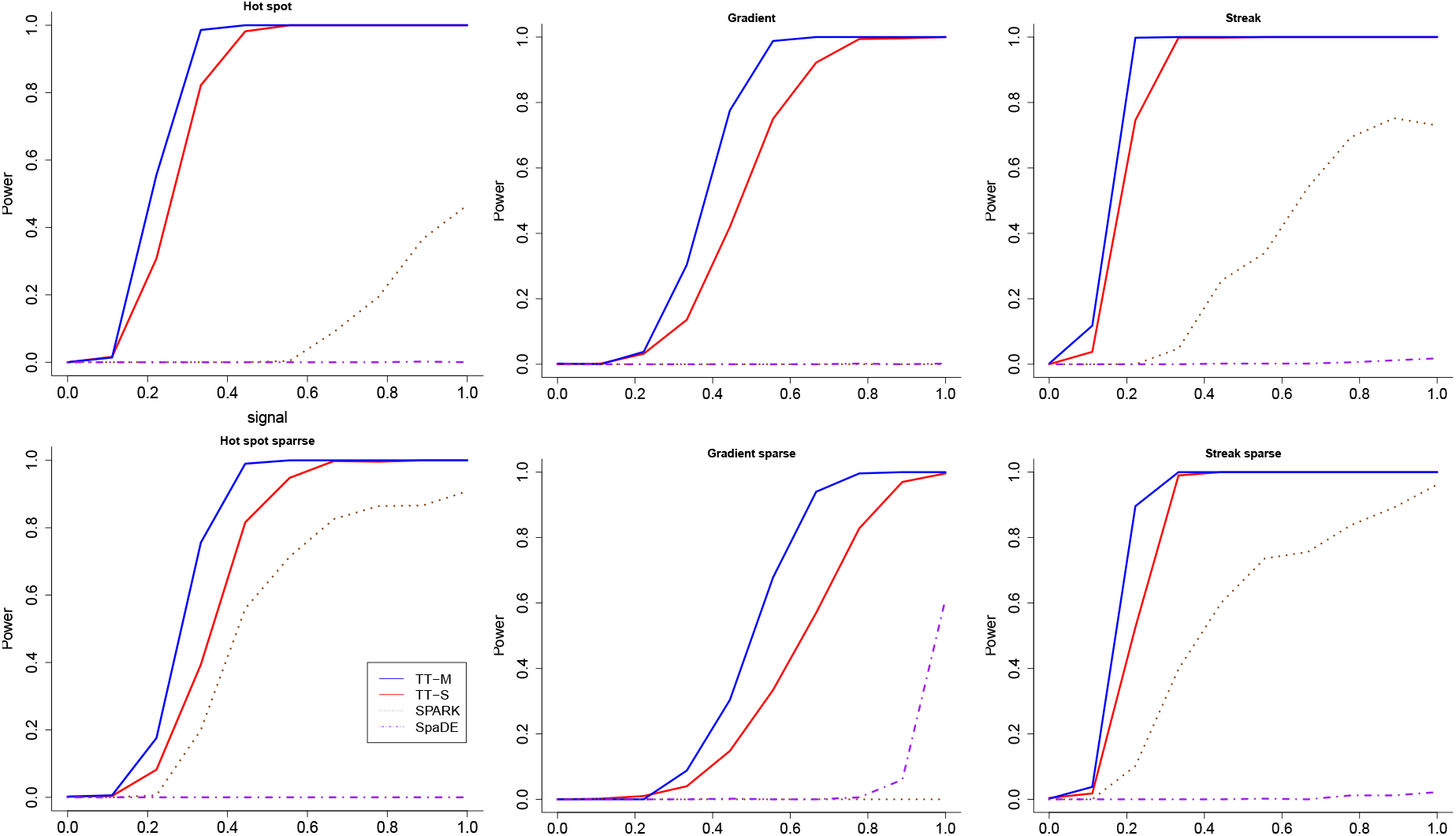
The power versus signal for the all the methods when *α* = 0.01. The first row are from the three patterns with non-sparse setting, and the second row are from the three patterns with sparse setting. The blue solid line and red solid line denote the multiple splits (TT-M) and single splits (TT-S). The green dashed line denote SPARK, while the purple dotted line denotes SpaDE, respectively.

### C Additional Real Data Analysis

#### C.1 Hypothalamus Data

In this section, we analyze a MERFISH dataset collected on the preoptic area of the mouse hypothalamus (Moffitt et al., 2018). The dataset contains 160 genes measured in 4,975 single cells. In the original study, 155 of the 160 genes were either labeled as markers of distinct cell populations or are relevant to various neuronal functions of the hypothalamus. Thus a large number of genes might be selected as spatially variable genes. We report the analysis result in Fig.14. As we expected, the empirical distribution of the *p*-values for our proposed methods are valid under the null condition, since their distributions are clearly below the diagonal line, see Fig.14.(a). On the other hand, SpaDE also produces uniformly valid *p*-values, while SPARK does not. The empirical distribution of *p*-values obtained by SPARK crosses the diagonal line several times, which indicates that the test could have a high false positive rate under the alternative hypothesis. Indeed, for the real data analysis, TT-S found 105 genes and TT-M found 115 genes, while SPARK found 143 genes and SpaDE found 124 genes. The genes detected by TT-S and TT-M all overlap with the findings of SPARK and SpaDE, which confirms the validity of the proposed methods. The GO annotation reveals that the detected genes are not only related to protein bindings, but also other cell functions such as transcription coactivator activity.

**Figure 14:**
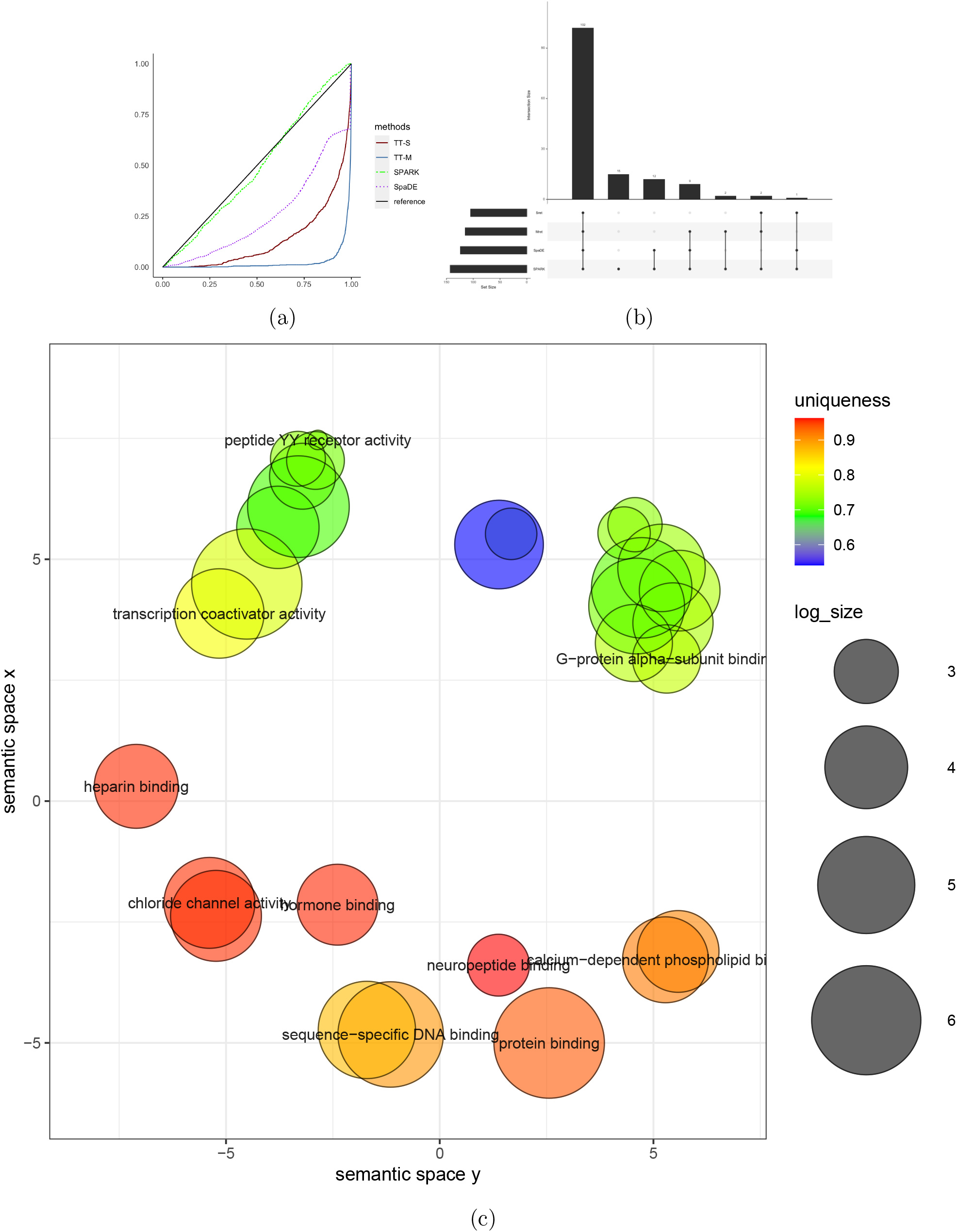
Analysis for the MERFISH dataset. (a): The empirical distribution of the *p*-values under the null condition in the permuted data. The blue solid line and red solid line denote the multiple splits (TT-M) and single splits (TT-S)2. 5The green dashed line denote SPARK, while the purple dotted line denotes SpaDE, respectively. (b): The upset plot shows the overlap of genes for TT-S and TT-M compared with SPARK and SpaDE. (c): The clustering of GO annotations for the genes detected by TT-M.

In terms of variable importance, %IncMSE equals 0.581 for the horizontal axis, 0.549 for the vertical axis, and is 0.665 for the interaction effect of the horizontal and vertical axis. Thus the spatial patterns are more equally spread across both the vertical and horizontal axis. This fact can also be observed from the 8 genes with the smallest *p*-values detected by TT-M, as illustrated in Fig.15.

**Figure 15:**
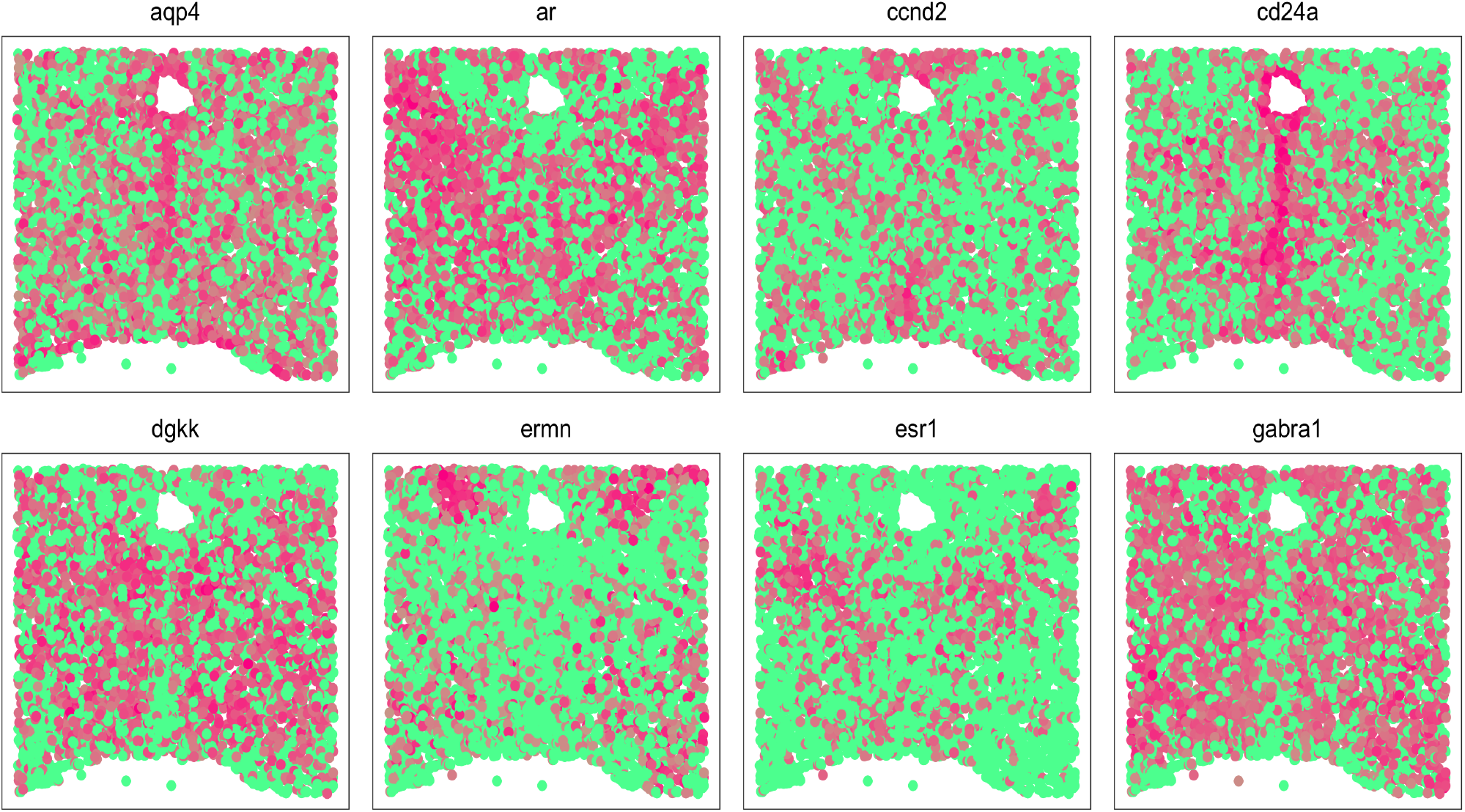
The 8 genes that have the smallest *p*-values detected by TT-M in the mouse hypothalamus data .

#### C.2 Hippocampus Data

The last dataset was a small seqFISH dataset with 249 genes measured on 131 single cells in the mouse hippocampus (Shah et al., 2016). SPARK found 17 genes and SpaDE found 11 genes. The results are reported in Figure 16. As expected, all methods produce valid *p*-values under the null condition. For the real analysis, TT-S did not find any genes while TT-M identified 3 genes (*lyve, mog, myl14*, shown in Fig.16c) and all of them overlap with the previous two approaches.

**Figure 16:**
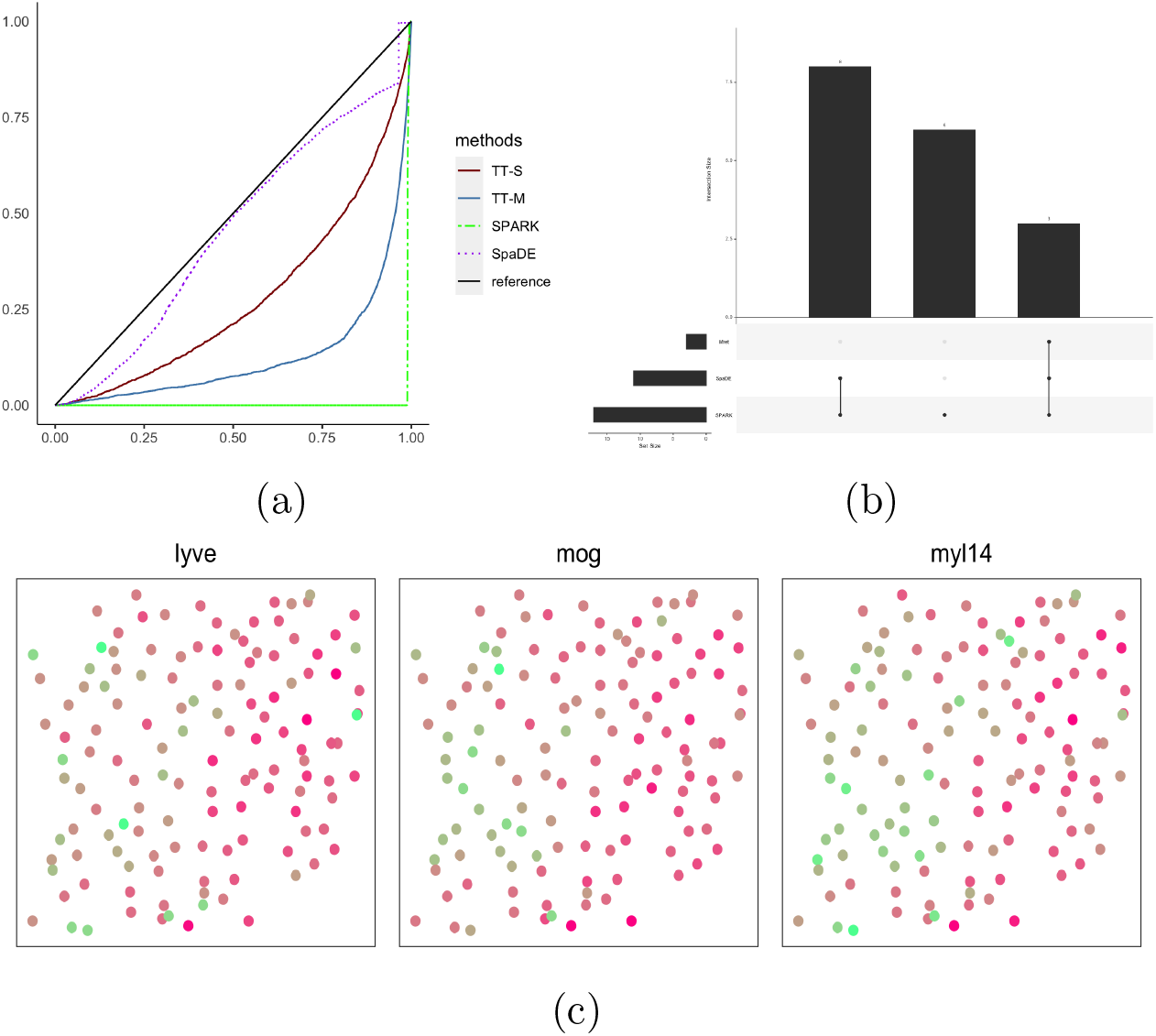
Analysis for the seqFISH dataset. (a): The empirical distribution of the *p*-values under the null condition in the permuted data. The blue solid line and red solid line denote the multiple splits (TT-M) and single splits (TT-S). The green dashed line denote SPARK, while the purple dotted line denotes SpaDE, respectively. (b): The upset plot shows the overlap of genes for TT-M compared with SPARK and SpaDE. (c): The 3 genes detected by TT-M in the mouse hippocampus data.

**Figure 17:**
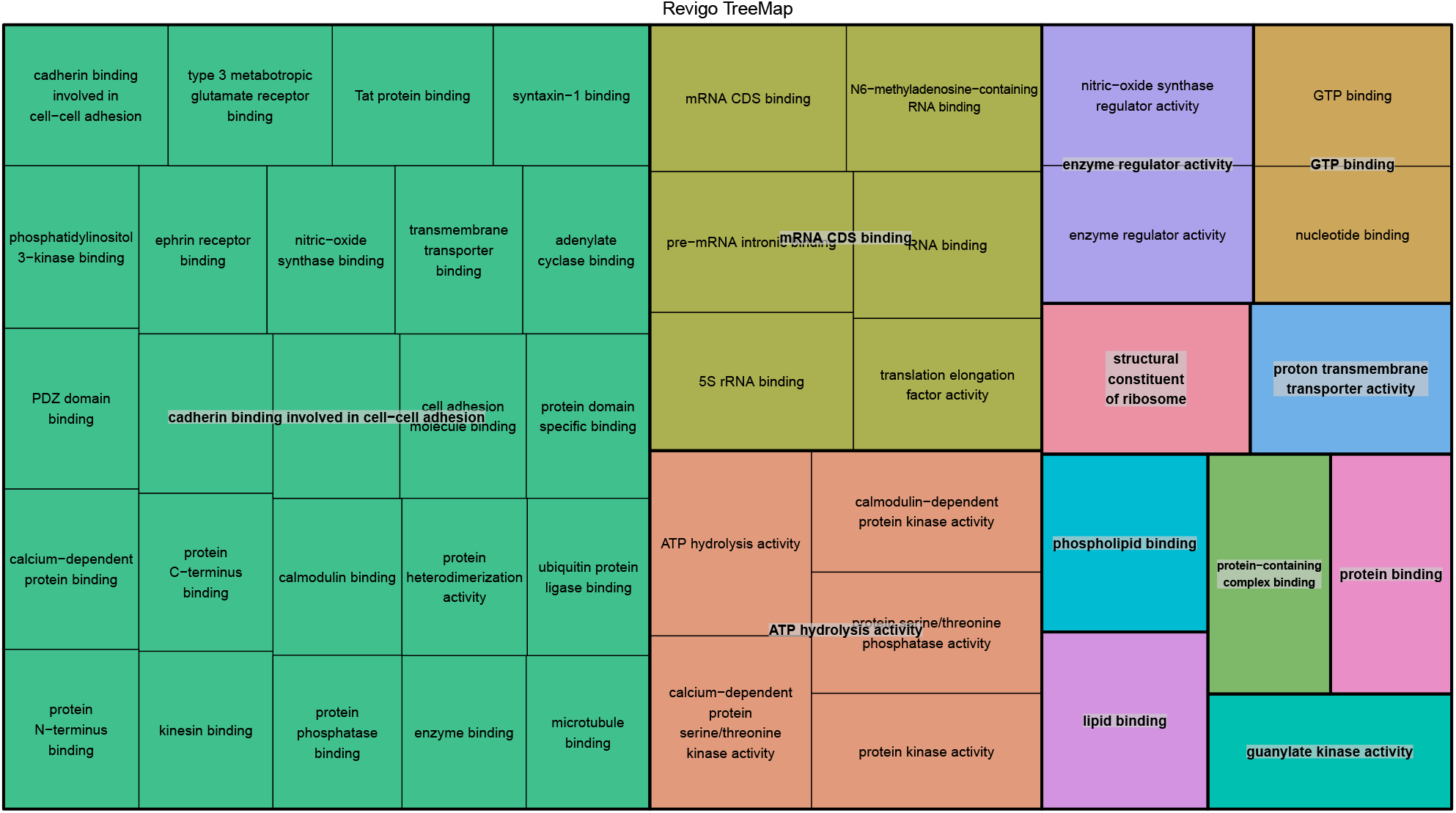
The mouse olfactory bulb data: Revigo tree map for the GO annotations based on genes detected by TT-S.

**Figure 18:**
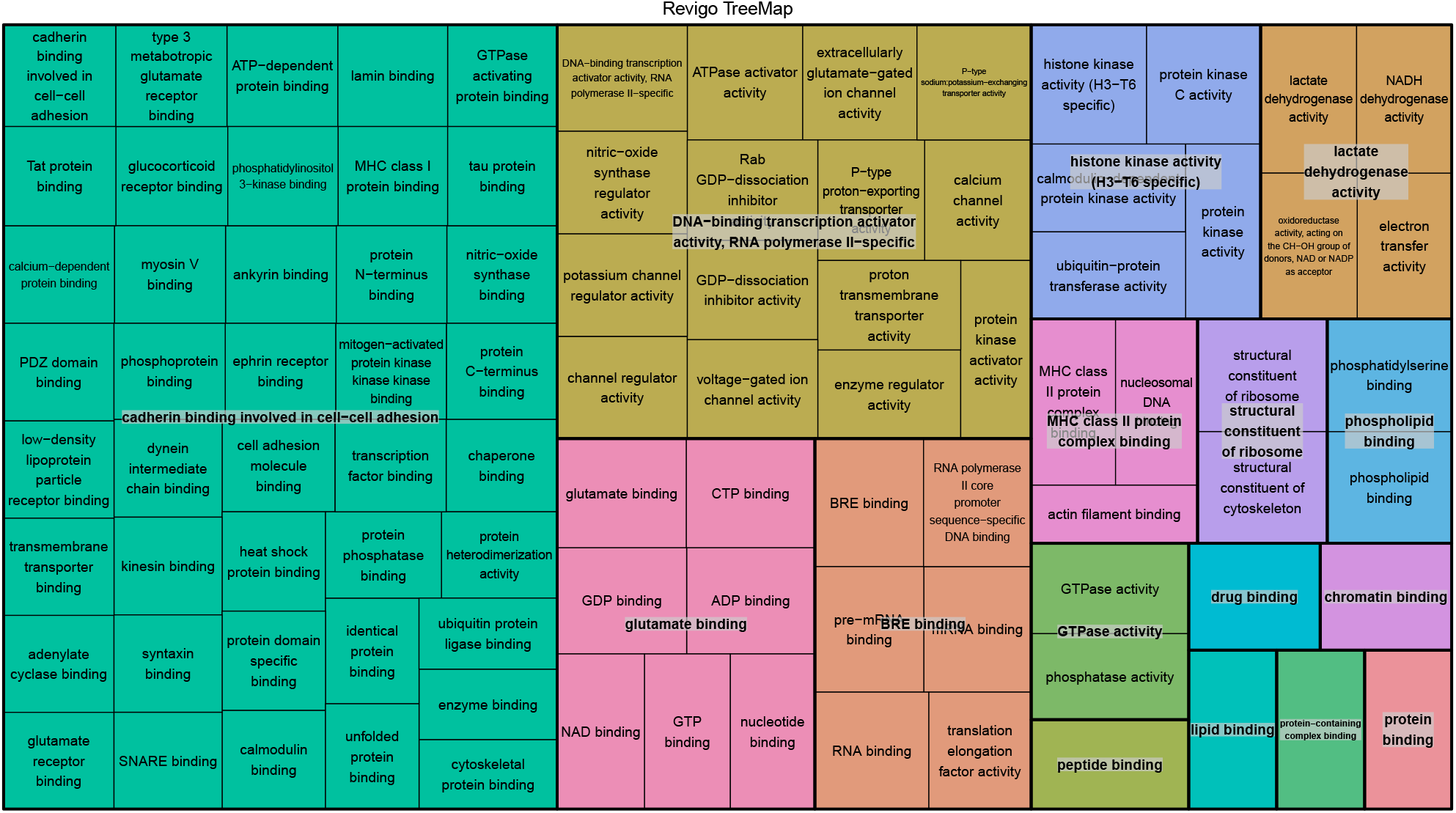
The mouse olfactory bulb data: Revigo tree map for the GO annotations based on genes detected by TT-M.

**Figure 19:**
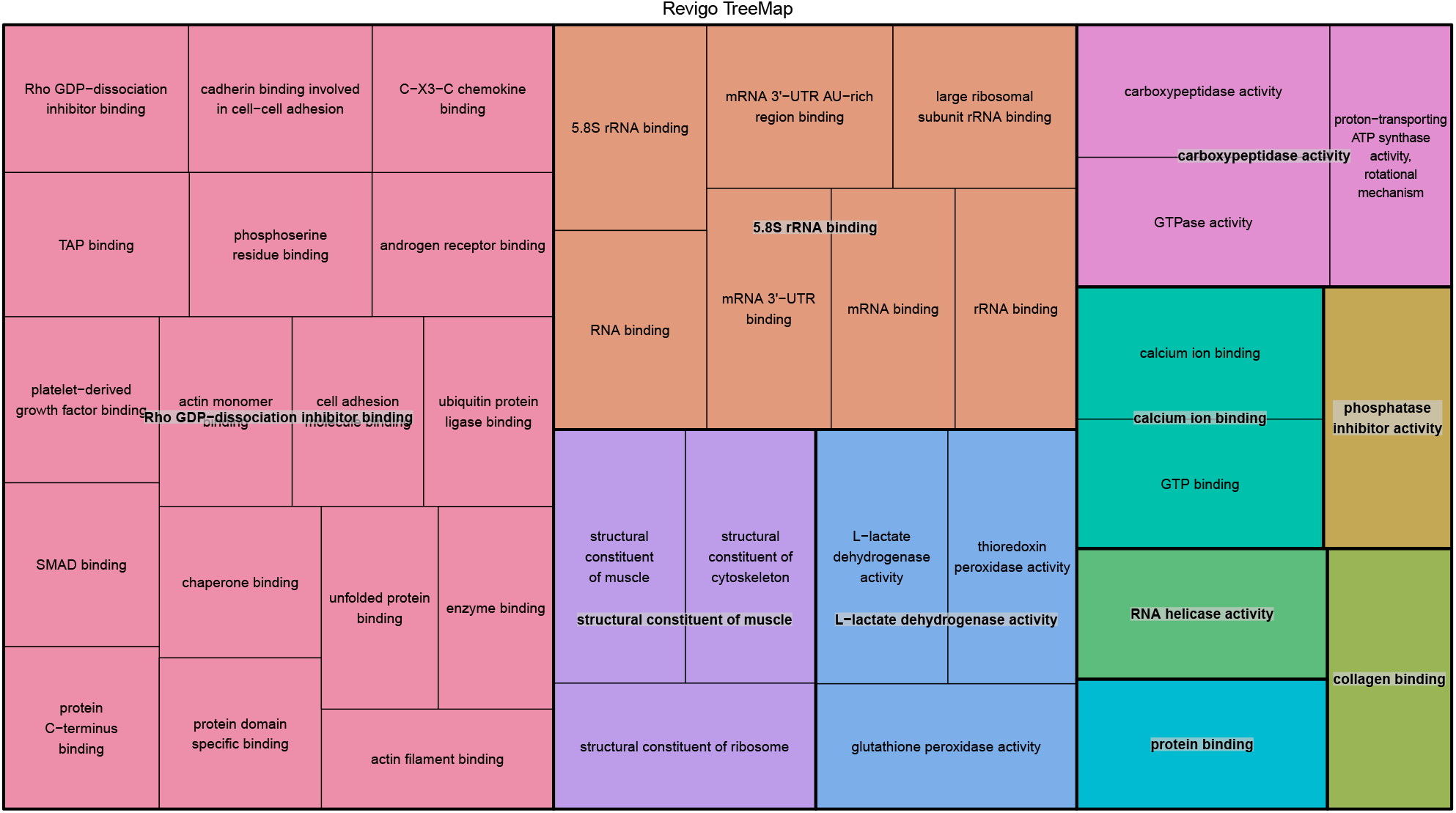
The breast cancer data: Revigo tree map for the GO annotations based on genes detected by TT-S.

**Figure 20:**
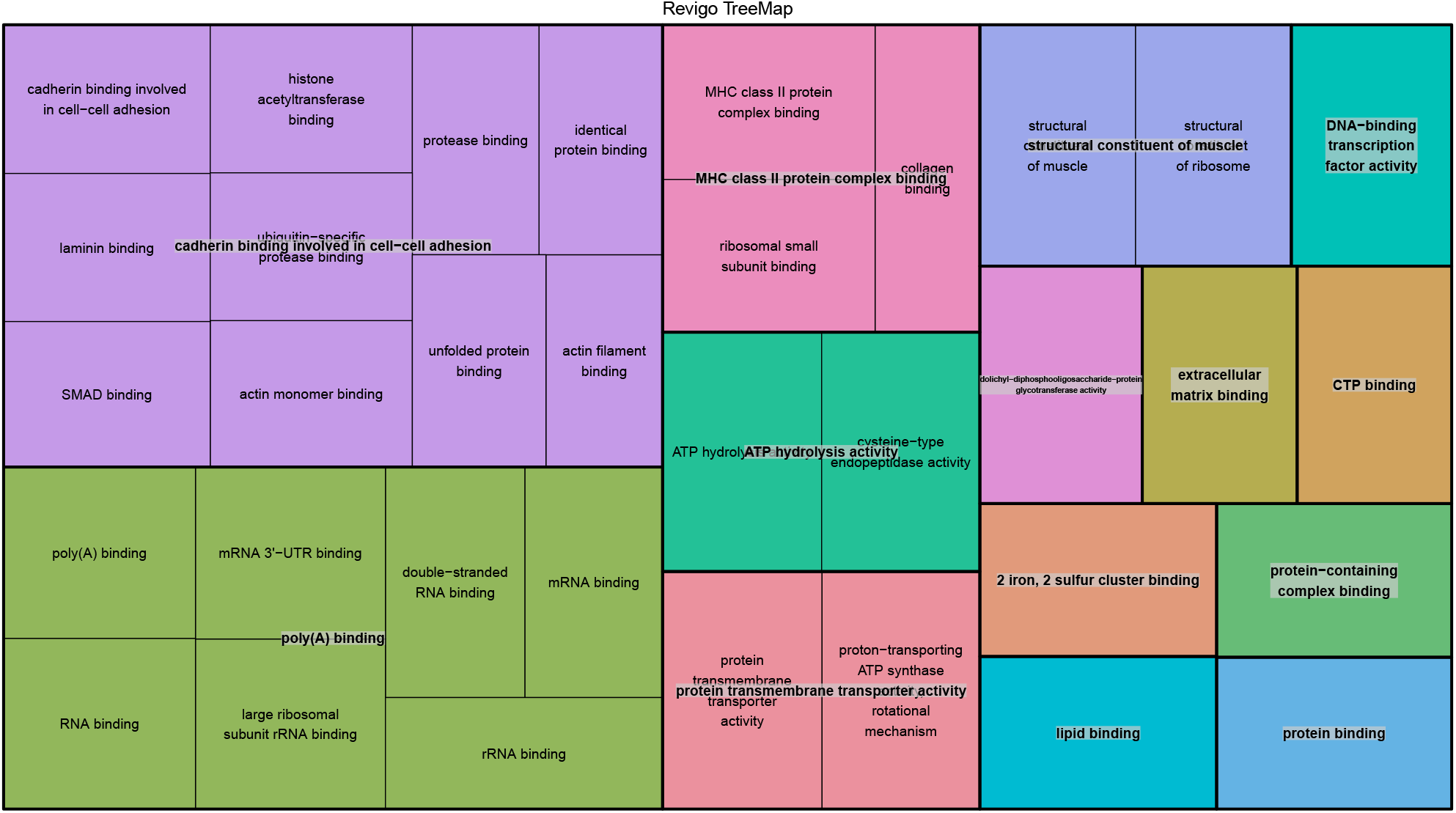
The breast cancer data: Revigo tree map for the GO annotations based on genes detected by TT-M.

**Figure 21:**
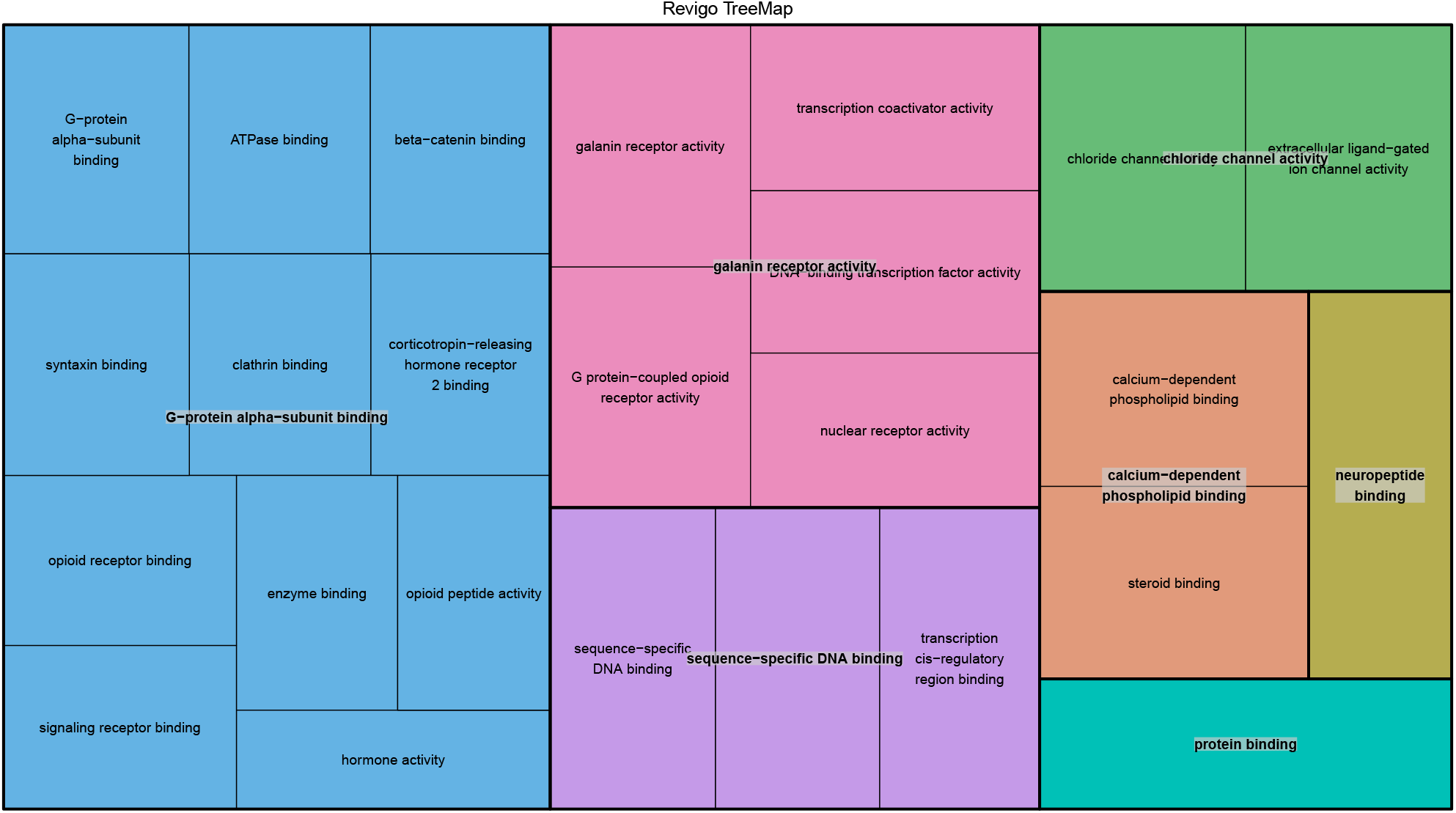
The mouse hypothalamus data: Revigo tree map for the GO annotations based on genes detected by TT-S.

**Figure 22:**
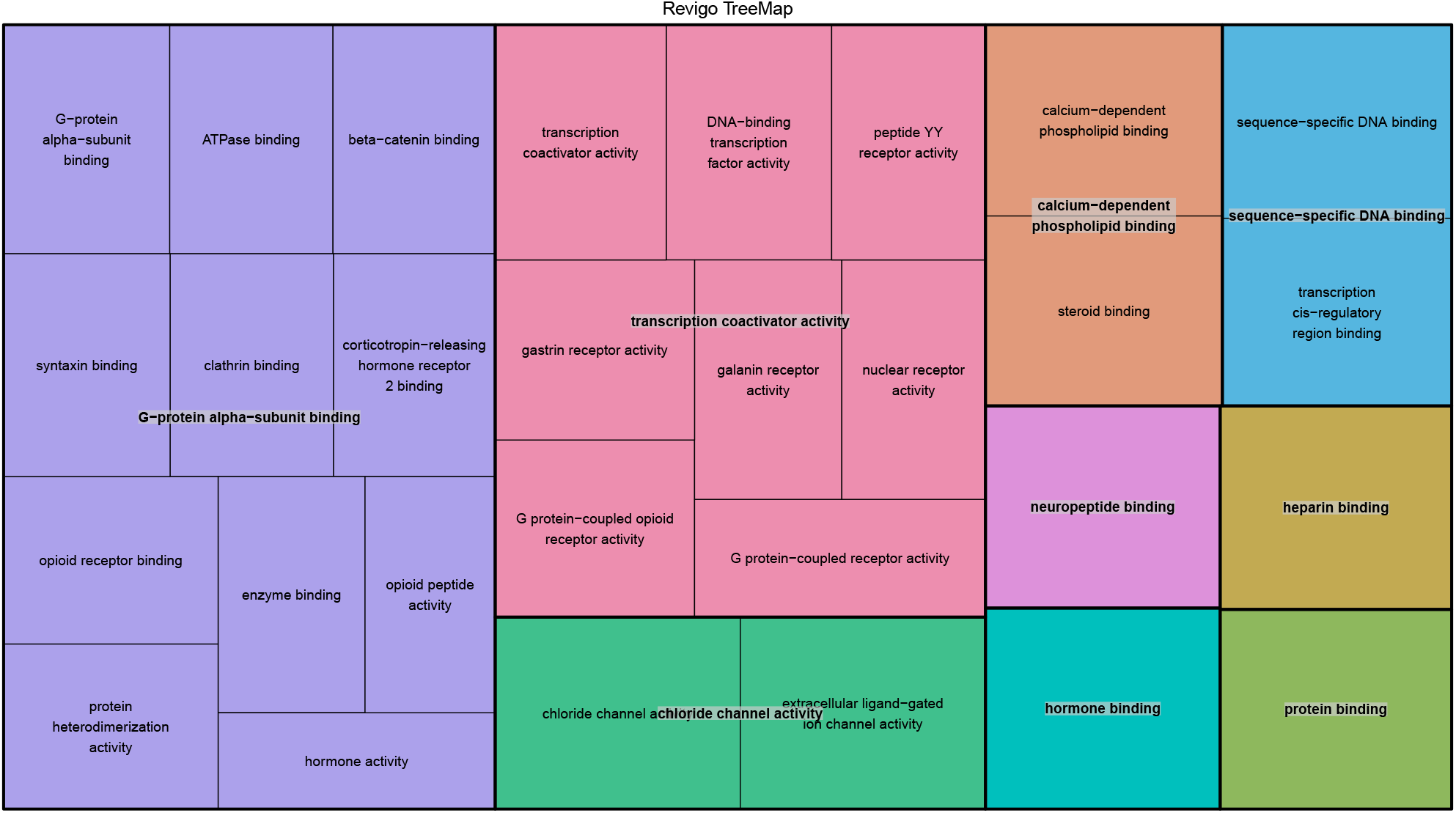
The mouse hypothalamus data: Revigo tree map for the GO annotations based on genes detected by TT-M.

**Figure 23:**
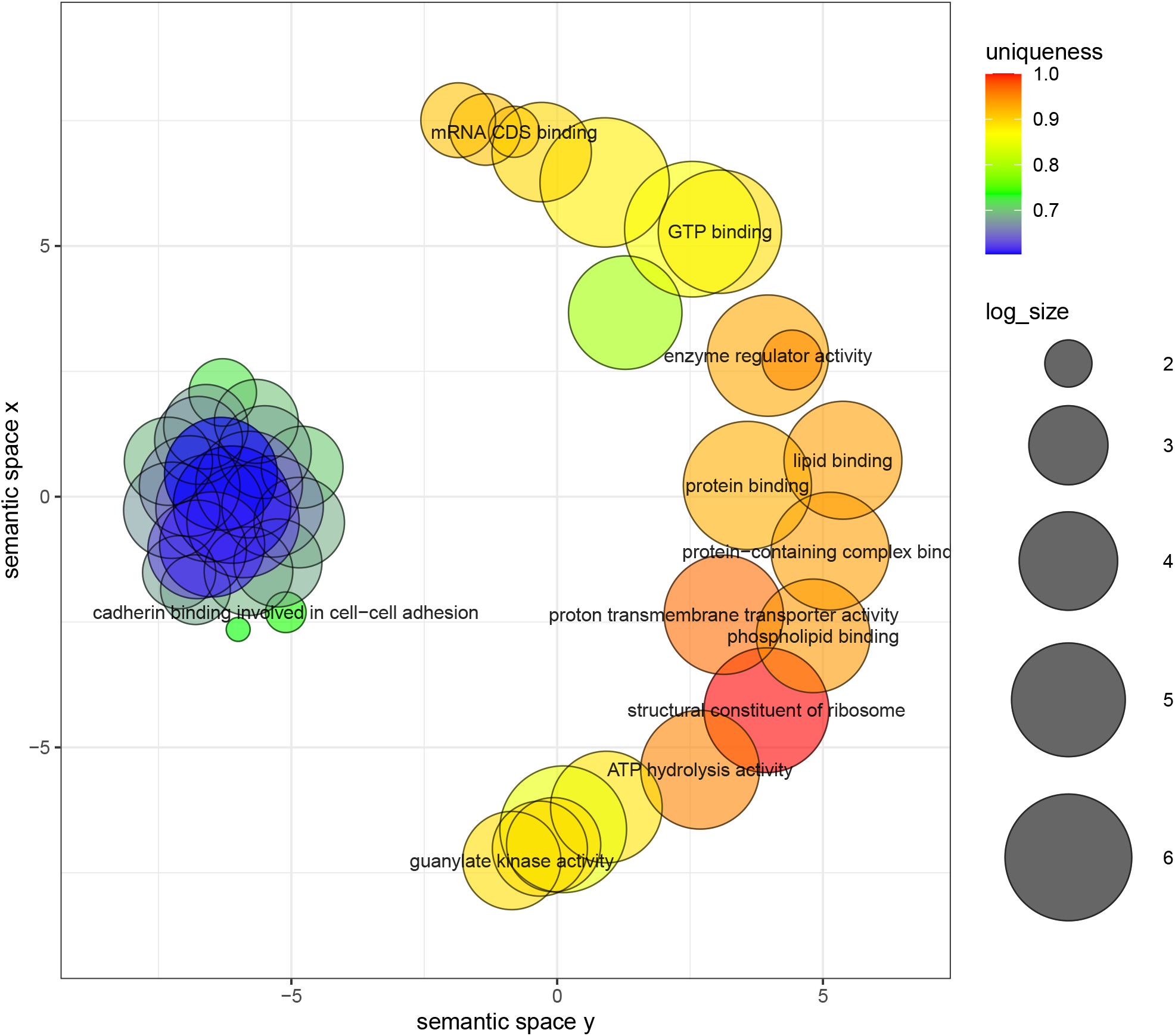
The mouse olfactory bulb data: clustering of GO annotations for the genes detected by TT-S.

**Figure 24:**
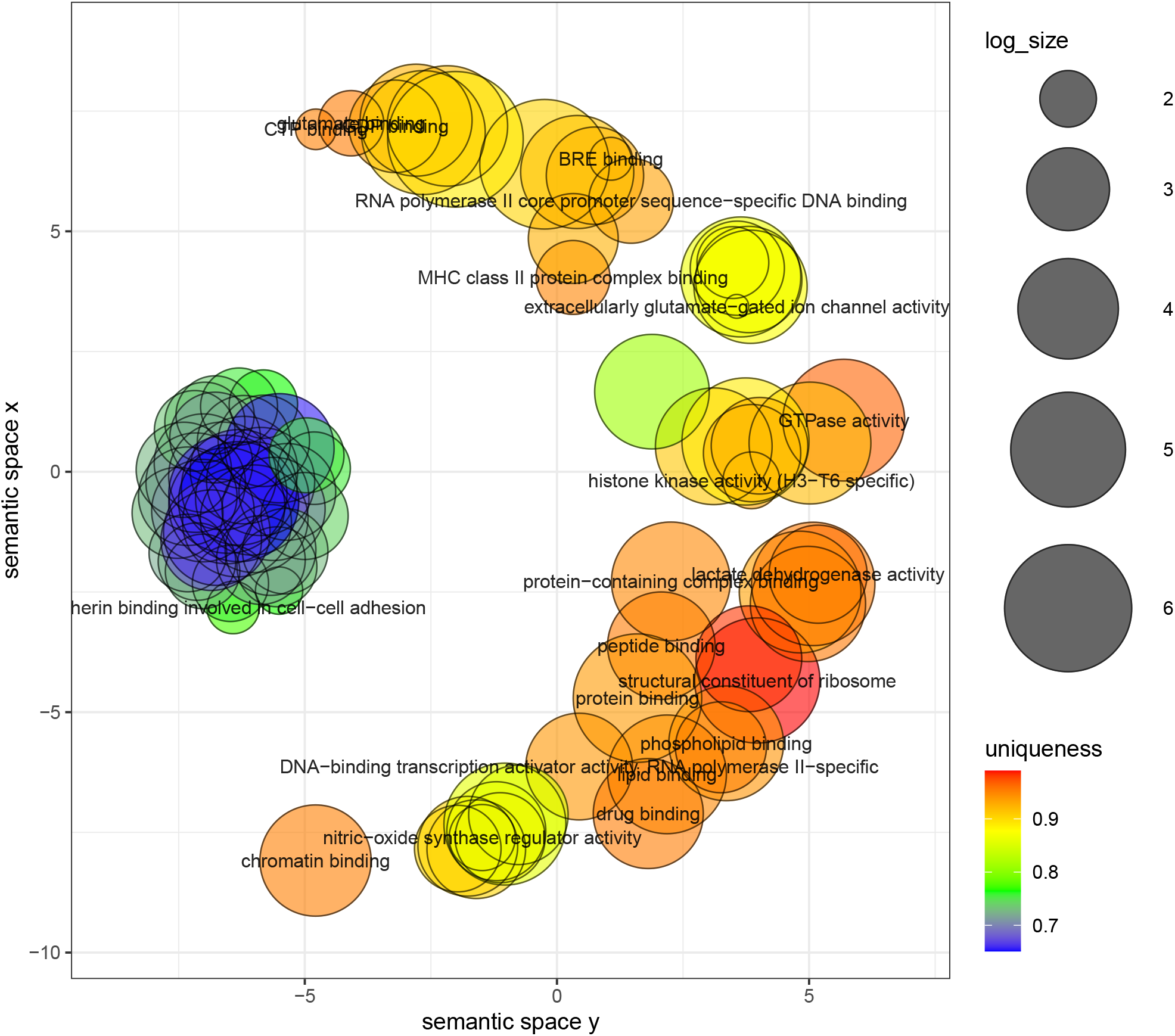
The mouse olfactory bulb data: clustering of GO annotations for the genes detected by TT-M.

**Figure 25:**
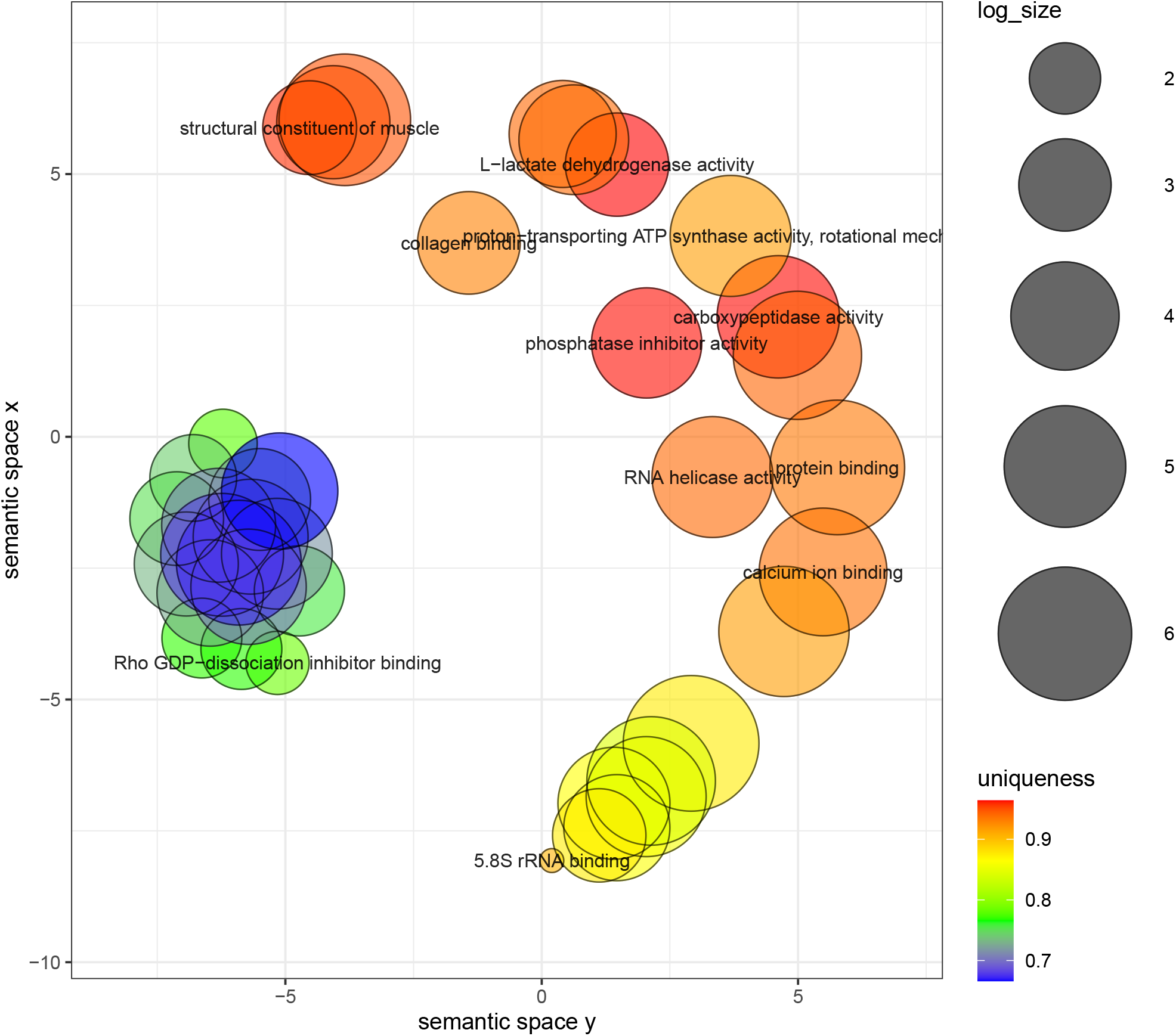
The breast cancer bulb data: clustering of GO annotations for the genes detected by TT-S.

**Figure 26:**
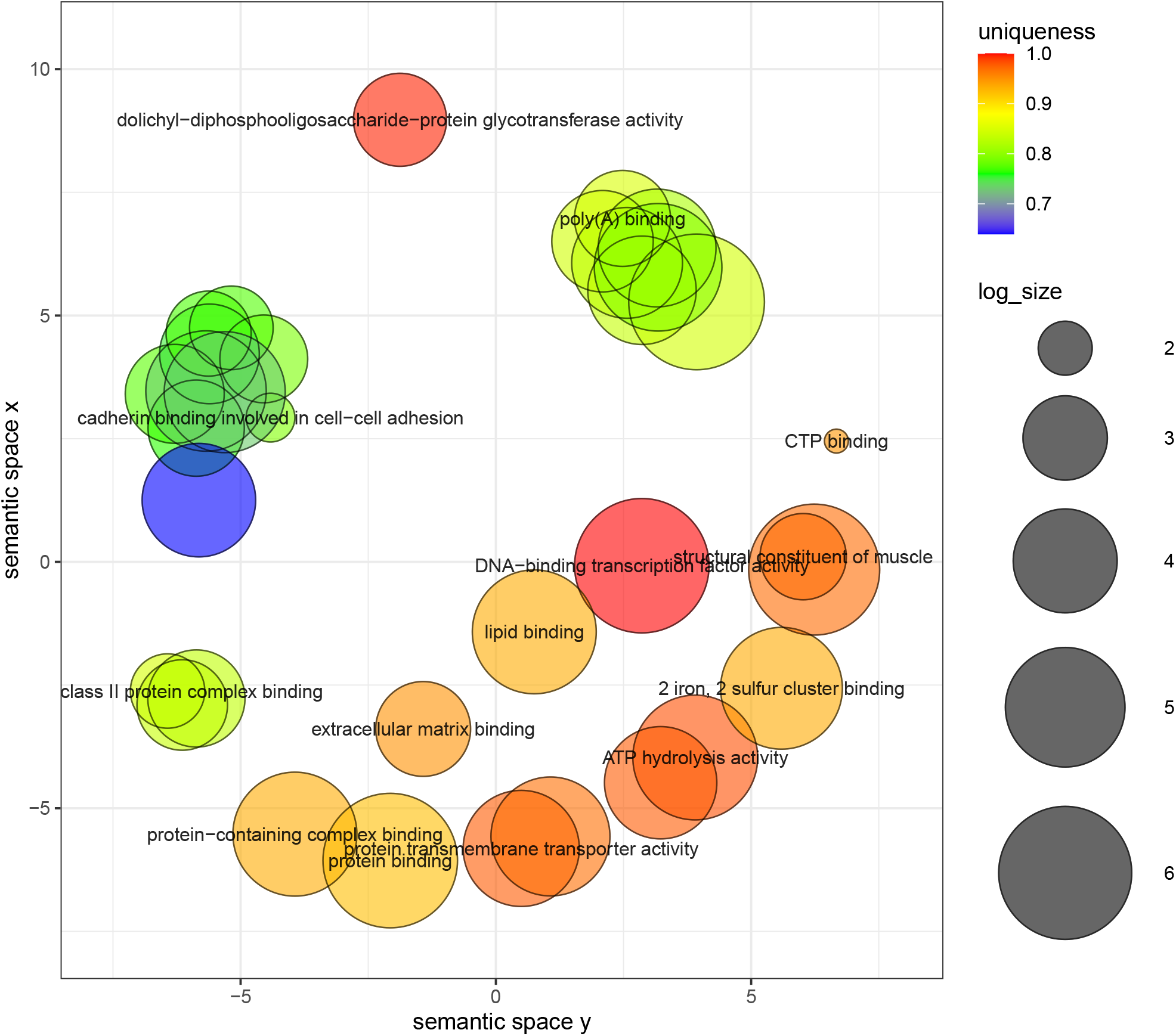
The breast cancer data: clustering of GO annotations for the genes detected by TT-M.

**Figure 27:**
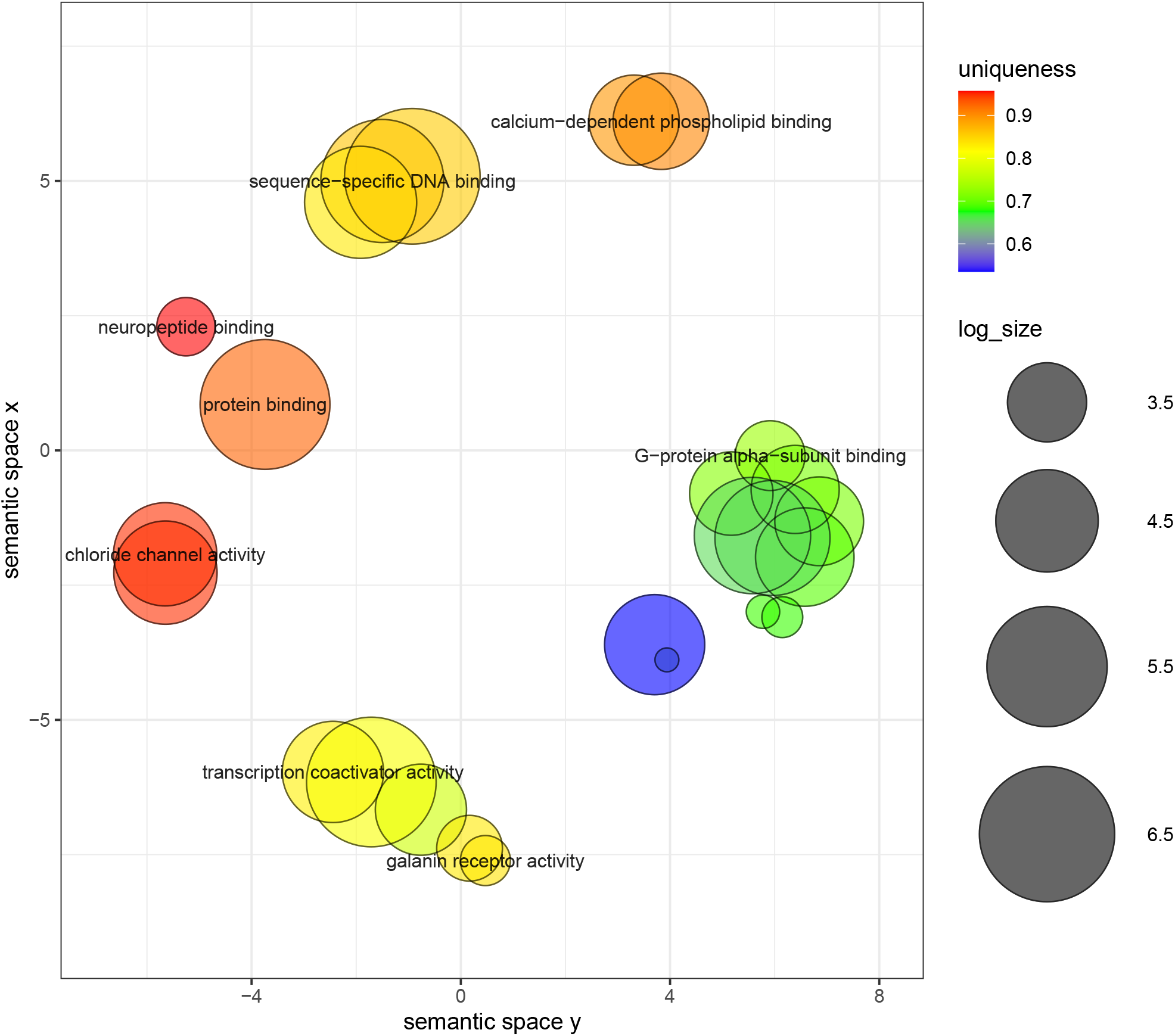
The mouse hypothalamus data: clustering of GO annotations for the genes detected by TT-S.

**Figure 28:**
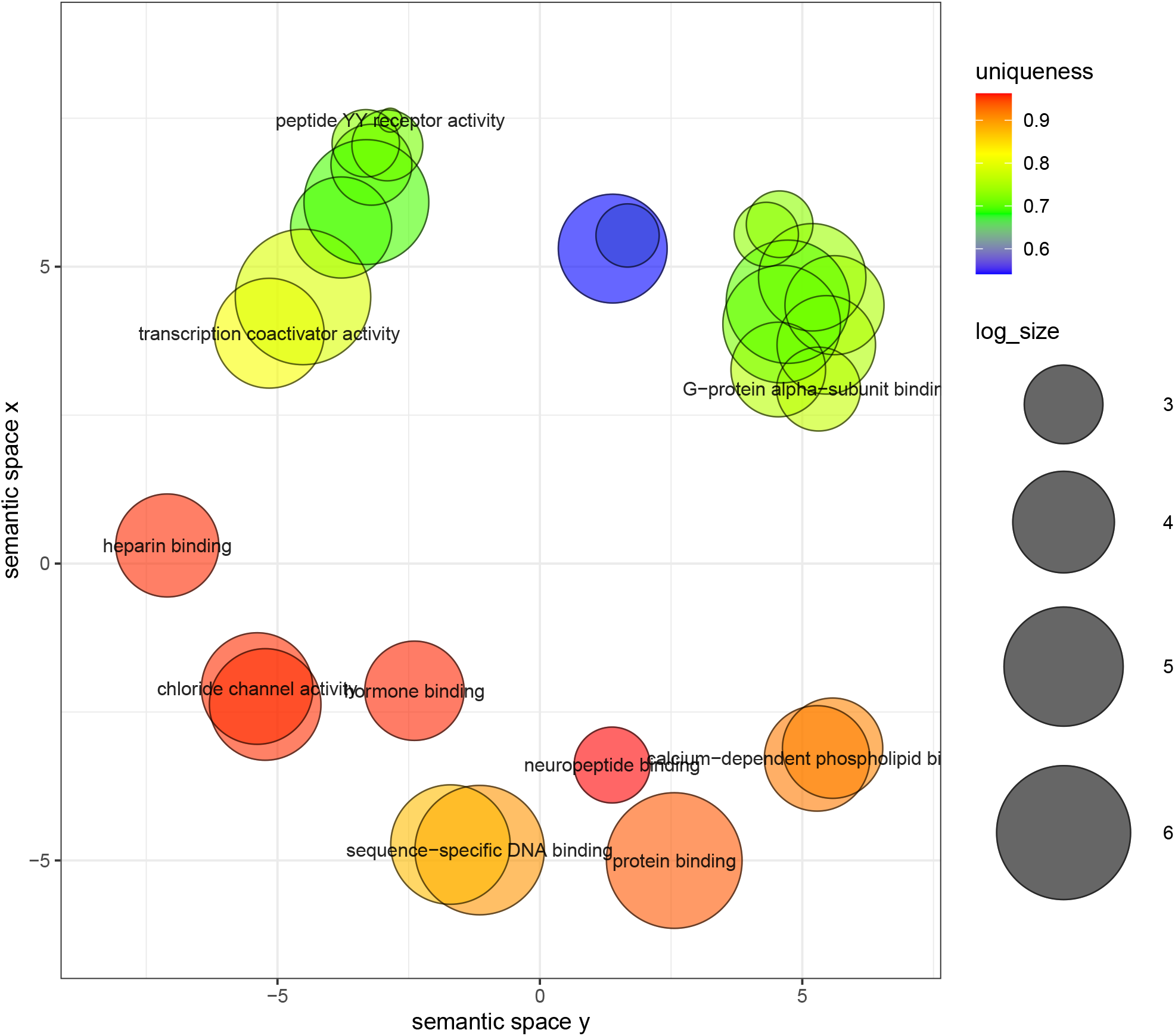
The mouse hypothalamus data: clustering of GO annotations for the genes detected by TT-M.

**Figure 29:**
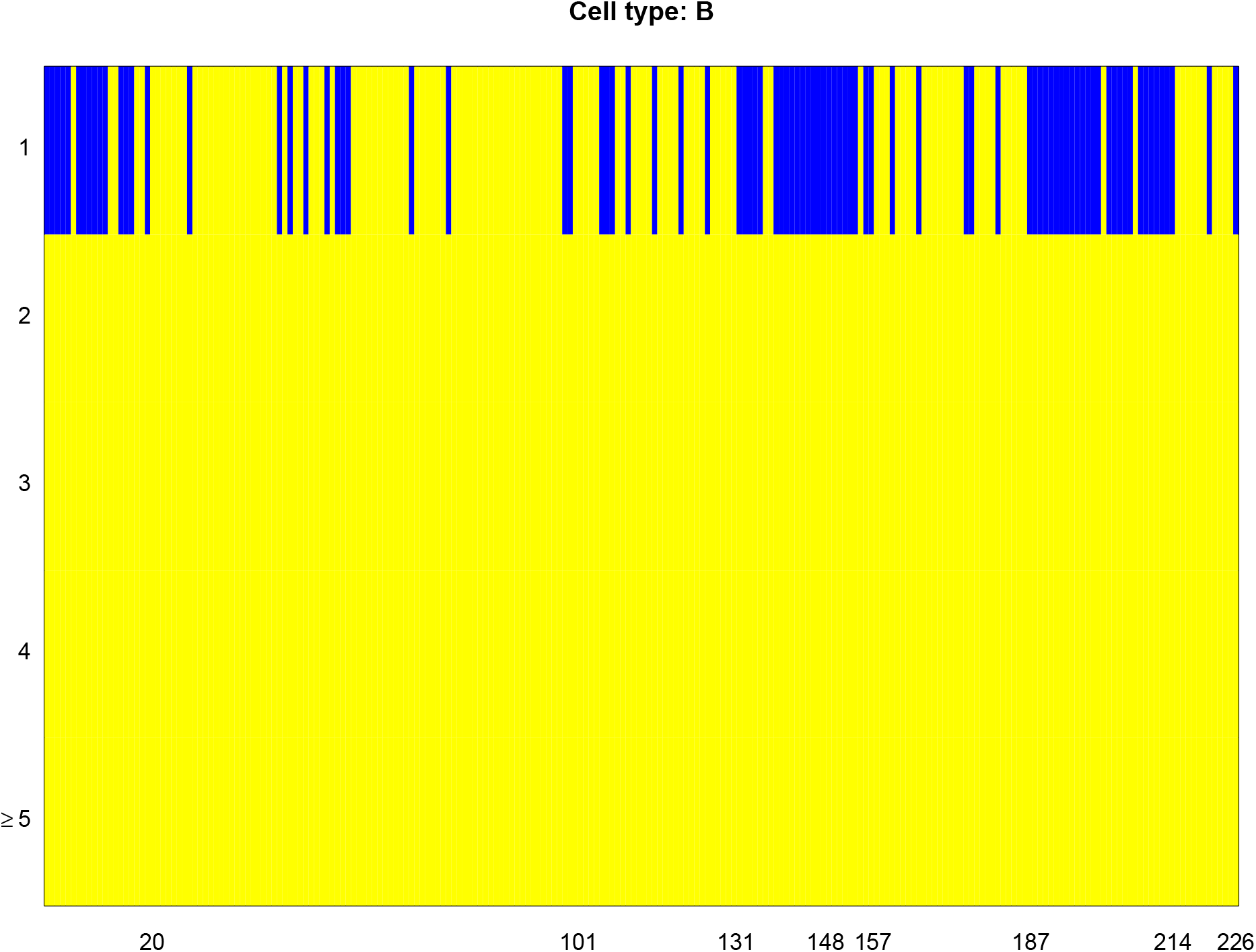
The predictability test of every proteins for B cells.

**Figure 30:**
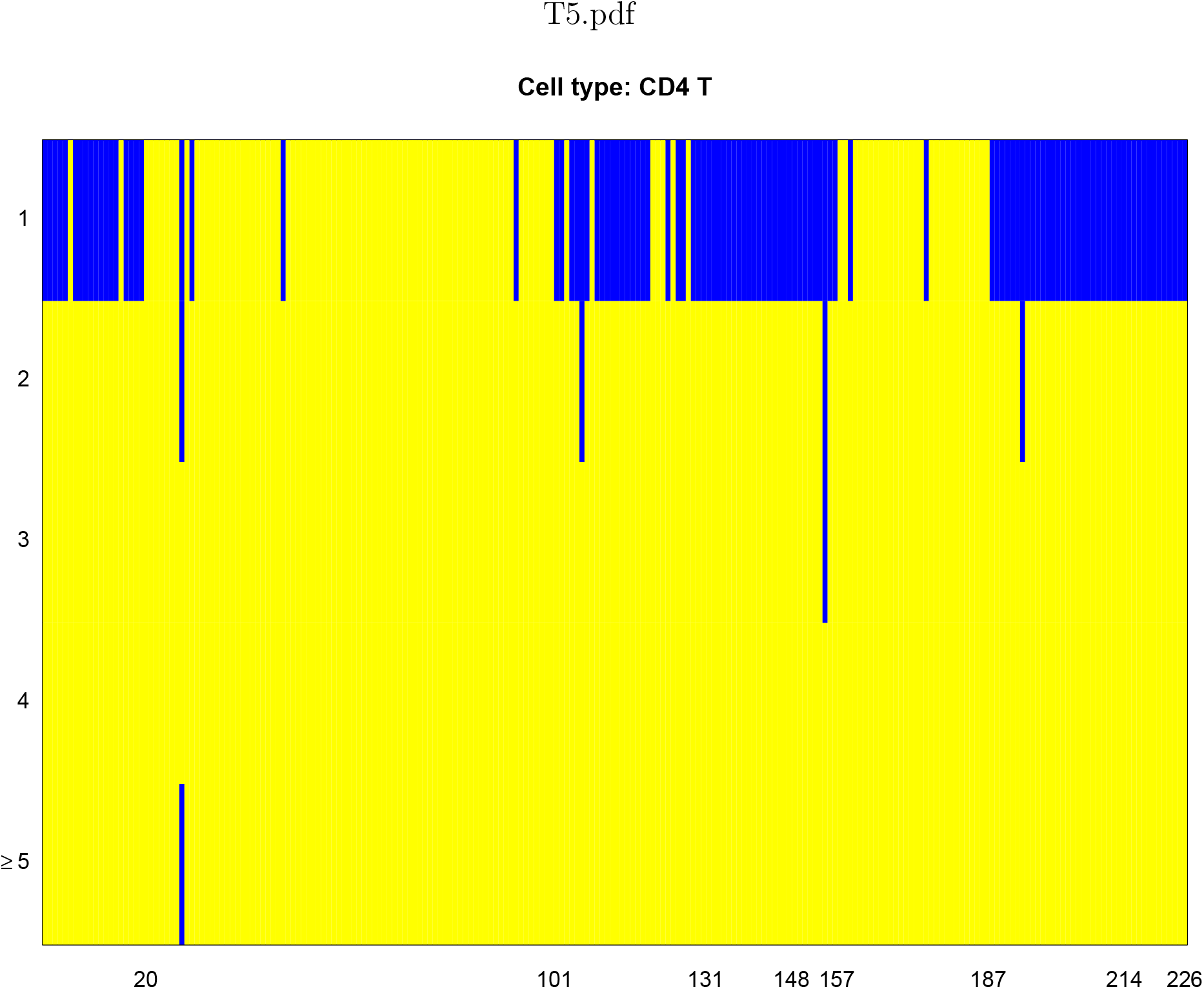
The predictability test of every proteins for CD4 cells.

**Figure 31:**
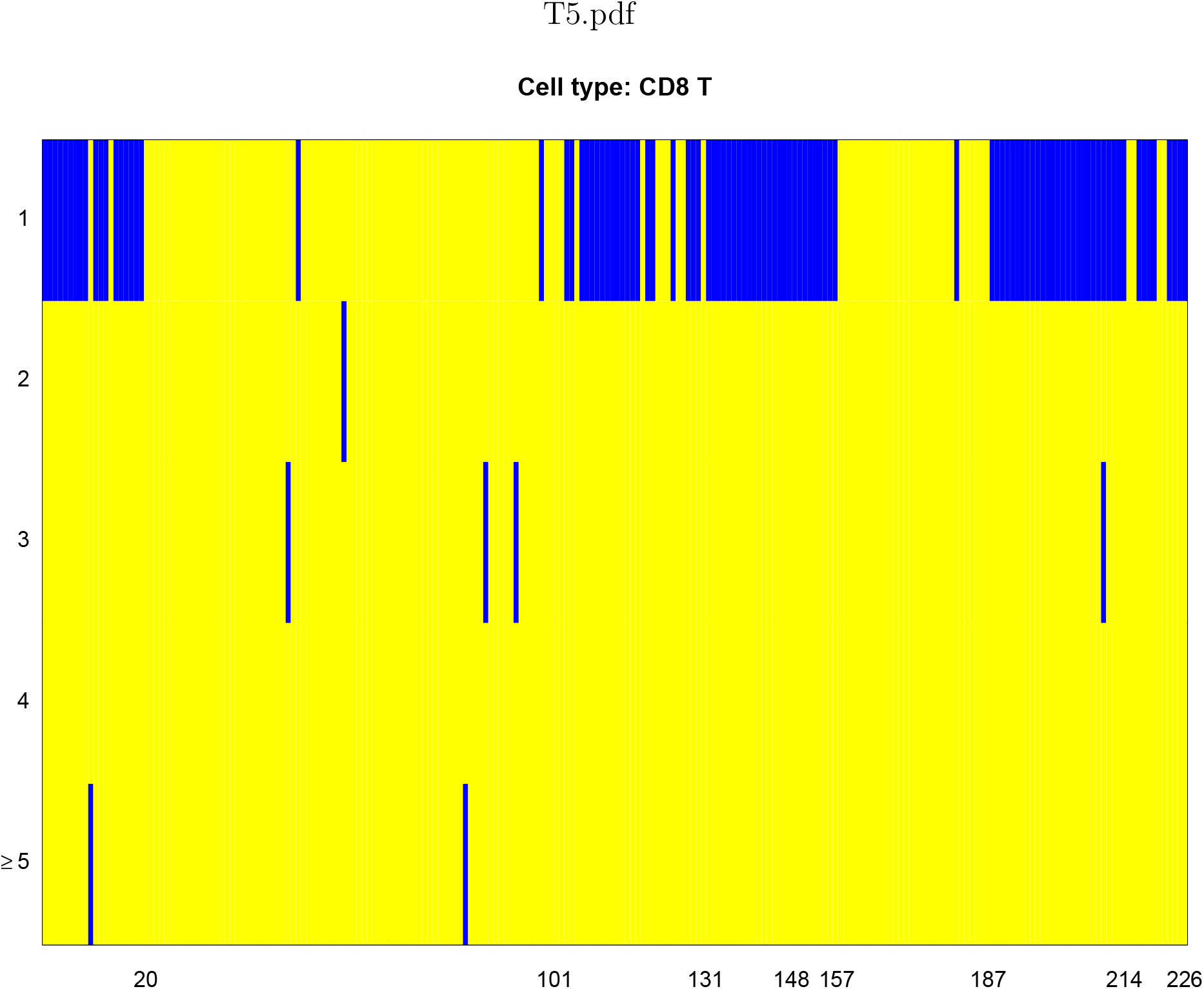
The predictability test of every proteins for CD8 T cells.

**Figure 32:**
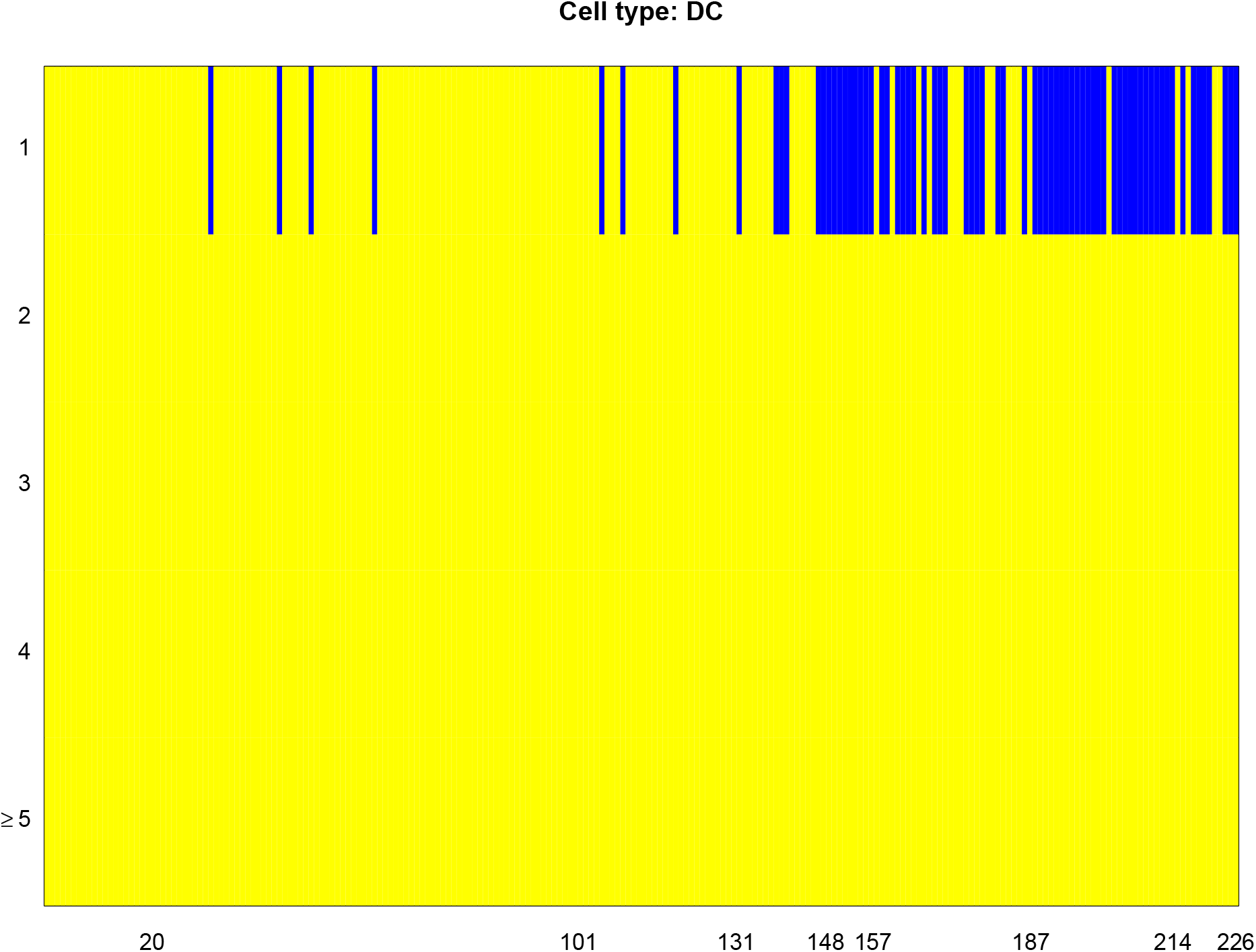
The predictability test of every proteins for DC cells.

**Figure 33:**
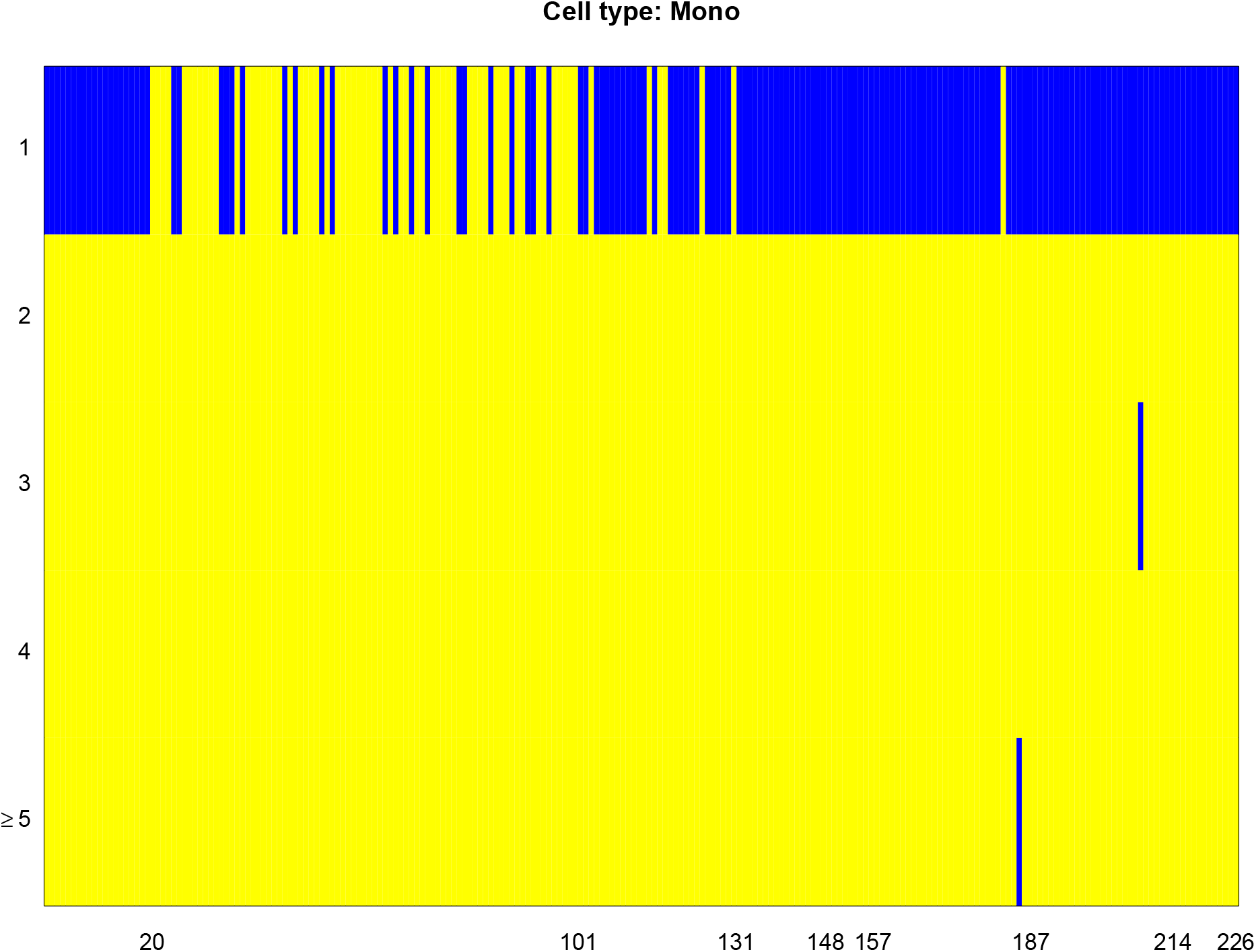
The predictability test of every proteins for Mono cells.

**Figure 34:**
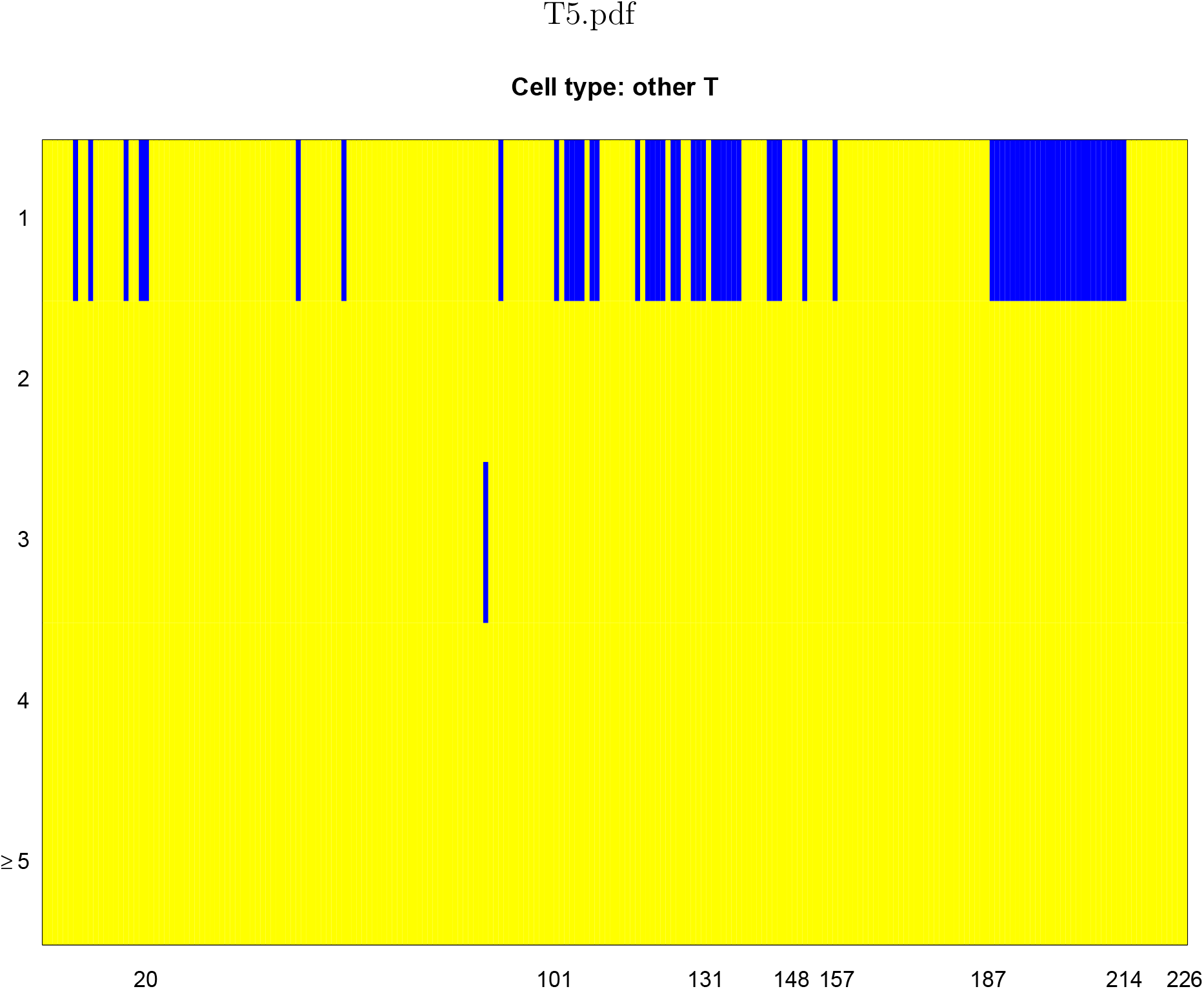
The predictability test of every proteins for other T cells.

**Figure 35:**
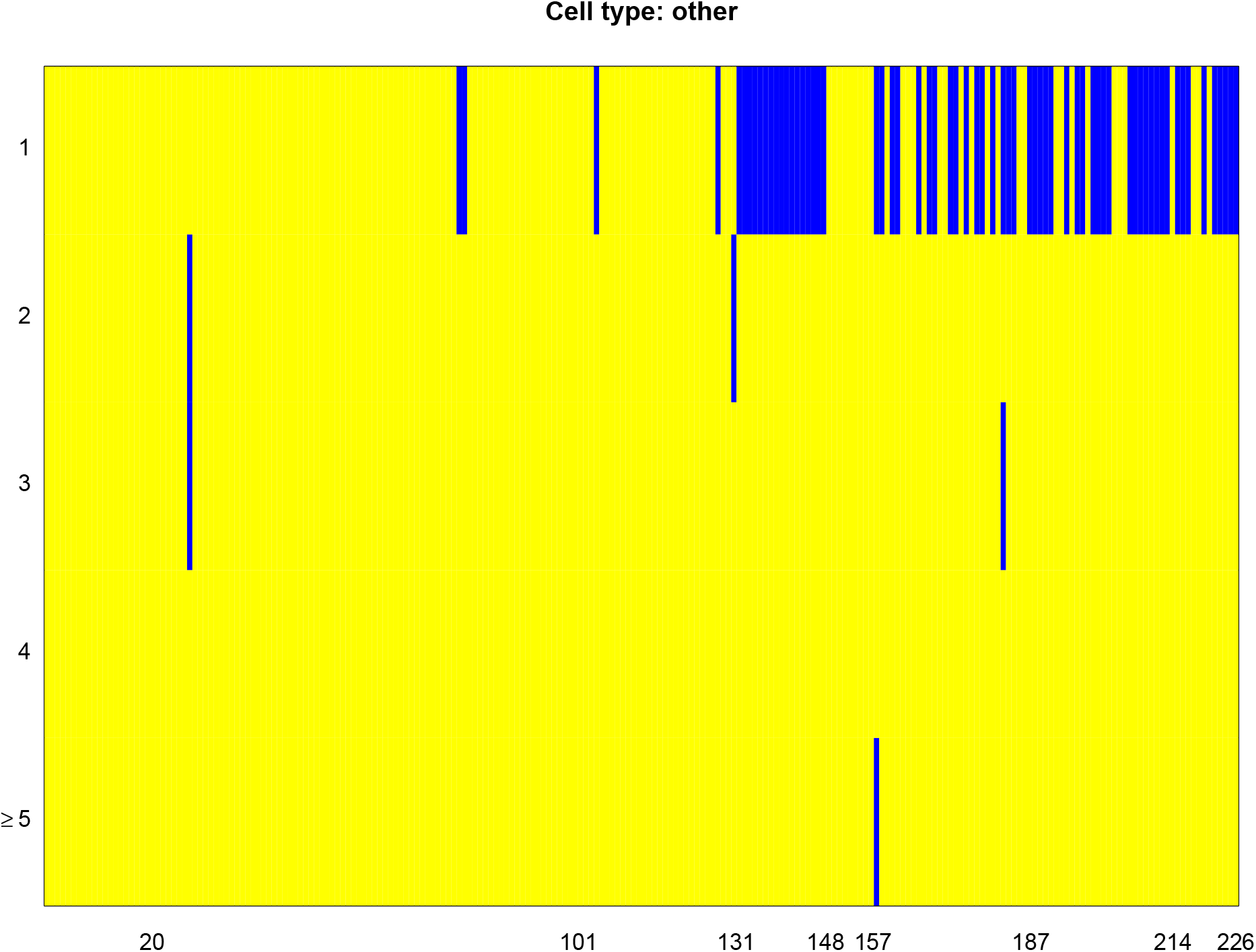
The predictability test of every proteins for other cells.

While all methods find less than 20 spatially variable genes for this dataset, we suspect the main reason that our method identifies the least amount of genes is that the sample size of 131 is too small for our method. The proposed method is based on sample splitting, thus the effective sample size for both regression and testing is at most 66 in this case. These samples are far from enough to train a proper machine learning model. As expected, we did not perform as well as the existing parametric models (SPARK or SpaDE).

## Details of Datasets

All datasets used in this paper are publicly available. We list the web source for each dataset below:

**CITE-seq Human PBMC data** https://atlas.fredhutch.org/data/nygc/multimodal/pbmc_multimodal.h5seurat

**Mouse olfactory bulb data** https://www.spatialresearch.org/resources-published-datasets/doi-10-1126science-aaf2403/

**Human breast cancer data** https://www.spatialresearch.org/resources-published-datasets/doi-10-1126science-aaf2403/

**Hypothalamus Data** https://datadryad.org/stash/dataset/doi:10.5061/dryad.8t8s248

**Hippocampus Data** https://www.cell.com/cms/10.1016/j.neuron.2016.10.001/attachment/759be4dc-04a6-4a58-b6f6-9b52be2802db/mmc6.xlsx

## Notes

### Competing Interest Statement

The authors have declared no competing interest.

